# Metal binding site alignment enables network-driven discovery of recurrent geometries across sequence-divergent proteins and drug off-targets

**DOI:** 10.64898/2026.03.05.709885

**Authors:** Vetle Simensen, Eivind Almaas

## Abstract

Metal-binding proteins account for nearly half of the characterized proteome, and they rely on metal-binding sites (MBSs) as critical determinants of their structural stability and biological function. However, methods for comparing their local binding environments lag behind those for whole-structure alignment. Here, we represent MBSs as atomic point clouds surrounding bound metal ligands and align them with a fine-tuned iterative closest point algorithm. Applying this framework to a redundancy-reduced collection of MBSs derived from all metalloproteins in the Protein Data Bank (PDB), we perform pairwise alignments across 23,342 sites to construct a similarity network of metal-binding environments. The resulting network topology recapitulates metal coordination chemistry and enzyme function: links are strongly enriched within metal types and across shared EC subclasses. Conserved metalloenzyme families form cohesive subnetworks; for example, the binuclear ureohydrolase domain appears as two tightly connected components that also capture atypical members such as the dinickel metformin hydrolase. We observe only a moderate global association between protein sequence and MBS geometry, yet many network links connect near-identical binding-site architectures across proteins with low sequence identity, consistent with either divergent evolution with local MBS conservation or candidate cases of molecular convergent evolution. Integrating network proximity with structural evidence of drug binding identifies drugs with enriched connectivity among their targets and predicts 528 drug–off-target combinations across 88 drugs and 151 human proteins, recovering both known off-targets (e.g., ADAM/ADAMTS for matrix metalloproteinase inhibitors) and proposing novel ones. The MBS network thus provides a scalable resource for probing metalloprotein evolution, functional convergence, and the structural basis of drug cross-reactivity.

**Author summary:** We study how metals shape protein structure and function by comparing metal-binding sites (MBSs) rather than whole proteins. We represent each MBS site as a point cloud of atoms surrounding the bound metal and align 23,342 sites from the Protein Data Bank (PDB) with a fine-tuned iterative closest point algorithm. This yields a similarity network whose links mirror metal coordination chemistry and enzymatic roles: sites binding the same metal or sharing enzyme classes cluster together, and conserved metalloenzyme families (e.g., binuclear ureohydrolases) form tight subnetworks that also capture atypical members such as a dinickel metformin hydrolase. Because highly similar MBS geometries often link proteins with low sequence identity, the MBS network highlights candidates consistent with either divergent evolution with locally conserved MBS architecture or convergent evolution toward similar coordination geometries in otherwise unrelated protein contexts. Overlaying known drug-binding sites lets us flag drugs whose targets are tightly connected and propose plausible off-targets, recovering known matrix metalloproteinase off-targets and suggesting new ones. Our approach offers a scalable map of metalloprotein relationships useful for studying evolution and anticipating drug cross-reactivity.

## Introduction

Metal-binding proteins are ubiquitous in biological systems, with about one-third of all known structures binding metal ions or metal-containing cofactors (1). These proteins serve a multitude of functions essential for cellular processes. Broadly, the bound metal play either of two roles: structural or functional. Structurally, the metal ligands are critical for the proper folding and stabilization of proteins into their native three-dimensional structure (2). Functionally, they frequently facilitate biochemical catalysis in metalloenzyme active sites, commonly serving as redox centers or conduits for electron transfer, or as structural scaffolds that orient substrates and stabilize transition states (3).

The functional activity of metal-binding proteins depends not only on the presence of the metal ligand but also on their co-localization with surrounding amino acid residues not directly involved in metal coordination. These residues contribute to the local electrostatic environment, constrain the geometry of the binding site, and mediate interactions with substrates or macromolecular partners. In metalloenzymes, such cooperative interactions are key to catalysis, with precisely arranged side chains working in tandem with the metal ion to promote biochemical transformation (4). Beyond catalysis, such metal–residue coupling also shapes the folding, allosteric regulation, and dynamic responsiveness of metal-binding domains across diverse protein families (5).

Because of its pivotal role in determining biological function, the regions of a gene encoding metal-binding sites (MBSs) exhibit a notable degree of evolutionary conservation, even amidst extensive sequence changes throughout the gene (6). Moreover, the amino acids comprising the MBSs are typically dispersed across the protein sequence and sometimes even interspersed among distinct polypeptides within oligomeric protein complexes (7). Only by the intricate folding of the polypeptide chains are the residues brought together into close proximity in a precise and arranged geometry (8). Identifying these MBS residues solely from the protein sequence is challenging for well-conserved proteins and nearly impossible for less conserved ones (9). Despite this, sequence alignment remains the prevailing approach for the functional annotation of proteins, relying on the idea that sequence similarity reflects structural resemblance and, consequently, functional similarity (10).

Since the rise of high-throughput structure determination and structural genomics in the 1990s, structural biology has advanced rapidly as a global endeavor to resolve the tertiary structures of proteins (11). Innovations in experimental procedures such as X-ray crystallography (12), nuclear magnetic resonance (NMR) spectroscopy (13), and cryo-electron microscopy (cryo-EM) (14), have greatly accelerated protein structure determination. The Protein Data Bank (PDB), the primary repository for experimentally resolved three-dimensional protein structures, now contains almost 250,000 entries (15). This wealth of structural data has fueled the development of powerful computational methods for protein comparison and alignment, enabling researchers to probe structure–function relationships (16) and, more recently, predict uncharacterized structures with remarkable accuracy using tools such as AlphaFold (17).

Despite these advances, most structural comparison methods are designed for global fold alignment and are ill-suited for analyzing localized regions such as MBSs. These methods identify optimal alignments that are based on a best fit of all atoms, residues or the polypeptide backbone, which overlooks the fact that MBSs are typically more structurally conserved or rigid compared to distal regions of the protein (18). Local site-comparison methods partially address this gap, including the sequence order-independent profile–profile alignment (SOIPPA) algorithm (19), a binding site similarity search and function (BSSF) approach (20), and TrixP; an index-based method for protein binding site comparison (21). Nevertheless, despite their computational efficiency and scalability, these approaches rely on a coarse site abstraction (e.g. C*α* representation), which blurs atom-level correspondences, making them unfit to account for detailed structural alignment of MBSs at the atomic level.

In their investigation of the origin of the oxygen-evolving complex (OEC) of photosystem II, Raymond et al. sought to identify broader structural and sequence homologs (22). To this end, they adopted a geometric approach based on the iterative closest point (ICP) algorithm, a method for aligning three-dimensional point clouds by iteratively minimizing distances between point correspondences (23). In their adaptation, MBSs were represented as point clouds defined by the Cartesian coordinates of atoms surrounding the metals. By applying ICP to pairs of MBSs, they identified rigid spatial transformations that minimized interatomic distances and revealed structural similarities between the MBS point clouds. Their ICP-based workflow recovered tight geometric matches between Mn-centered environments in the OEC and metal sites from other PDB proteins, illustrating how point-cloud registration can surface local coordination environments that are not obvious from sequence or fold comparisons.

Building on this foundation, we extend and formalize the ICP-based framework to compare every structurally characterized MBS in the PDB. We align fixed-size point clouds of atoms nearest each metal ligand with a fine-tuned ICP variant and use the resulting scores to assemble a global MBS network. Although the representation is deliberately simple, it yields robust geometric comparisons at scale and a network topology enriched for shared metal coordination chemistry and enzymatic function. Because network links mirror structural compatibility, the network also supports discovery: it highlights conserved motifs across protein families, recurrent MBS geometries across sequence-divergent proteins, and potential drug off-targets when overlaid with known drug-binding sites. Together, this resource connects local MBS geometry to biological function and pharmacological interaction, providing a practical framework for studying metalloprotein function, evolution, and drug specificity.

## Methods

### Dataset construction and extraction of metal-binding site point clouds

We obtain protein structures from the RCSB PDB via its REST API (accessed 2024-07-20) (24). We query the archive for entries containing at least one of the target metal chemical components listed in Table 1. Each metal instance is treated as a candidate MBS and is subject to the following quality filter: We retain only crystallographic structures with reported overall resolution *<* 3.0 Å. Coordinates are parsed from PDBx/mmCIF files and expanded into biological assemblies by applying the PDB-provided assembly generation operators using Biopython (25; 26). For entries with multiple biological assemblies, we process each assembly separately and track assembly identifiers to enable downstream de-duplication.

**Table 1.** Metals used to identify MBSs. Elements used in the PDB query to retrieve metal-containing protein structures forming the basis for MBS point cloud extraction.

| Element | Symbol |
| --- | --- |
| Cobalt | Co |
| Copper | Cu |
| Iron | Fe |
| Manganese | Mn |
| Molybdenum | Mo |
| Nickel | Ni |
| Tungsten | W |
| Vanadium | V |
| Zinc | Zn |

To define a fixed-size representation for subsequent geometric comparison, we first calibrate the typical number of nearby protein atoms in metal-centered neighborhoods. For a stratified random subset of metal-containing structures, we enumerate all *protein* atoms within a 7 Å radius of each retained metal-ion coordinate. This choice contrasts with more commonly used, smaller cutoffs (e.g., 5 Å), which are frequently employed to delineate MBSs, but which tend to emphasize mainly atoms directly involved in metal coordination (27). By adopting a larger cutoff, we include both first-shell coordination and second-shell/scaffold context which are critical for biological function. The empirical atom count is found to converge to an average of approximately 65 protein atoms, which is subsequently used as the fixed-size, operational definition of an MBS point cloud.

For each biological assembly, we construct an MBS point cloud for every metal instance. Point clouds are built by ranking all protein atoms by Euclidean distance to the metal coordinate and selecting the *N* = 65 nearest atoms. This procedure yields a fixed-size set while permitting a variable effective radius across sites. Only atoms belonging to protein polymer entities are eligible for selection. Non-protein heteroatoms (e.g., ligand-derived groups, coenzymes, and solvent molecules) are excluded from the point cloud by construction. The polymer-chain entity contributing the largest fraction of atoms to a given point cloud is used to assign protein-specific metadata. Enzyme Commission (EC) numbers are obtained from the PDB when available. For missing EC assignments, we supplement annotations by cross-reference mapping PDB polymer chains to UniProt accessions and retrieving EC information from UniProt (28). When multiple EC numbers map to a given chain, we retain the full set of candidate EC annotations and propagate this ambiguity rather than collapsing to a single label.

### Robust ICP alignment of MBS point clouds

We use the iterative closest point (ICP) algorithm (23) to perform pairwise alignment of MBS point clouds. ICP computes a rigid spatial transformation that minimizes the squared distance between corresponding points in two point clouds *P* = {**p**_1_, …, **p**_*m*_}and *Q* = {**q**_1_, …, **q**_*n*_}in ℝ^*d*^ (29). The point registration problem can be formulated as finding an optimal transformation (**R**^(∗)^, **t**^(∗)^) that produces a minimal alignment error *ξ*(*P, Q*):

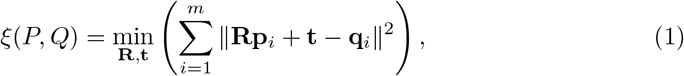

where **R** ∈ ℝ^*d×d*^ is a rotation matrix, **t** ∈ ℝ^*d*^ is a translation vector, and ∥ · ∥ is the Euclidean norm. The ICP algorithm iteratively solves this minimization problem by alternating between two steps:

- *Correspondence step*: Given the transformation (**R**^(*k*)^, **t**^(*k*)^) at iteration *k* of the algorithm, we determine the closest point 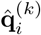 in *Q* for each point **p**_*i*_ ∈ **P** by minimizing the distance

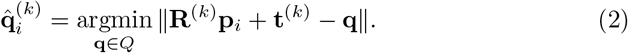
- *Alignment step*: We calculate the transformation matrices (**R**^(*k*+1)^, **t**^(*k*+1)^) of the (*k* + 1)th iteration by solving

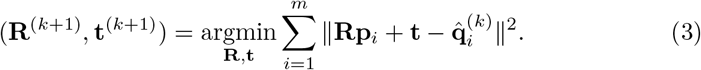

We employ the Open3D library for representing point clouds and conducting the ICP alignments (30). As residuals (i.e., the distances between point correspondences) are measured using squared distances (Eq.1), the metric is sensitive to outliers due to the disproportionate influence of poorly aligned points. To reduce this sensitivity, we incorporate Tukey’s biweight function (31) as a loss function.

The *correspondence* and the *alignment* steps are repeated until a convergence criterion is met, at which point we assume (**R**^(∗)^, **t**^(∗)^) are obtained. We employ two convergence criteria: (i) Limit the iteration count to *k* ≤ 30; or (ii) A relative change in root mean square deviation (RMSD) between successive iterations of *<* 10^−6^. The RMSD is defined in terms of the alignment error *ξ* as:

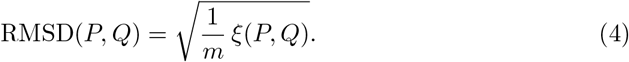

### Multi-start coarse-to-fine initialization for reliable ICP convergence

Because ICP optimizes a non-convex objective, it is only guaranteed to converge to a local minimum and is therefore sensitive to initialization (32). In preliminary experiments on MBS point clouds, repeated runs on the same geometrically similar pairs yield a characteristic bimodal distribution of final RMSD values (Fig 1 A), consistent with convergence either to a low-RMSD basin or to a higher-RMSD local minimum. Notably, the higher RMSD-mode overlap substantially with the RMSD distribution obtained from aligning unrelated point clouds with no meaningful geometric resemblance (S1 Fig in S1 File), motivating an explicit multi-start strategy to increase the probability of reaching the best-observed solution for a given pair.

**Fig 1.**
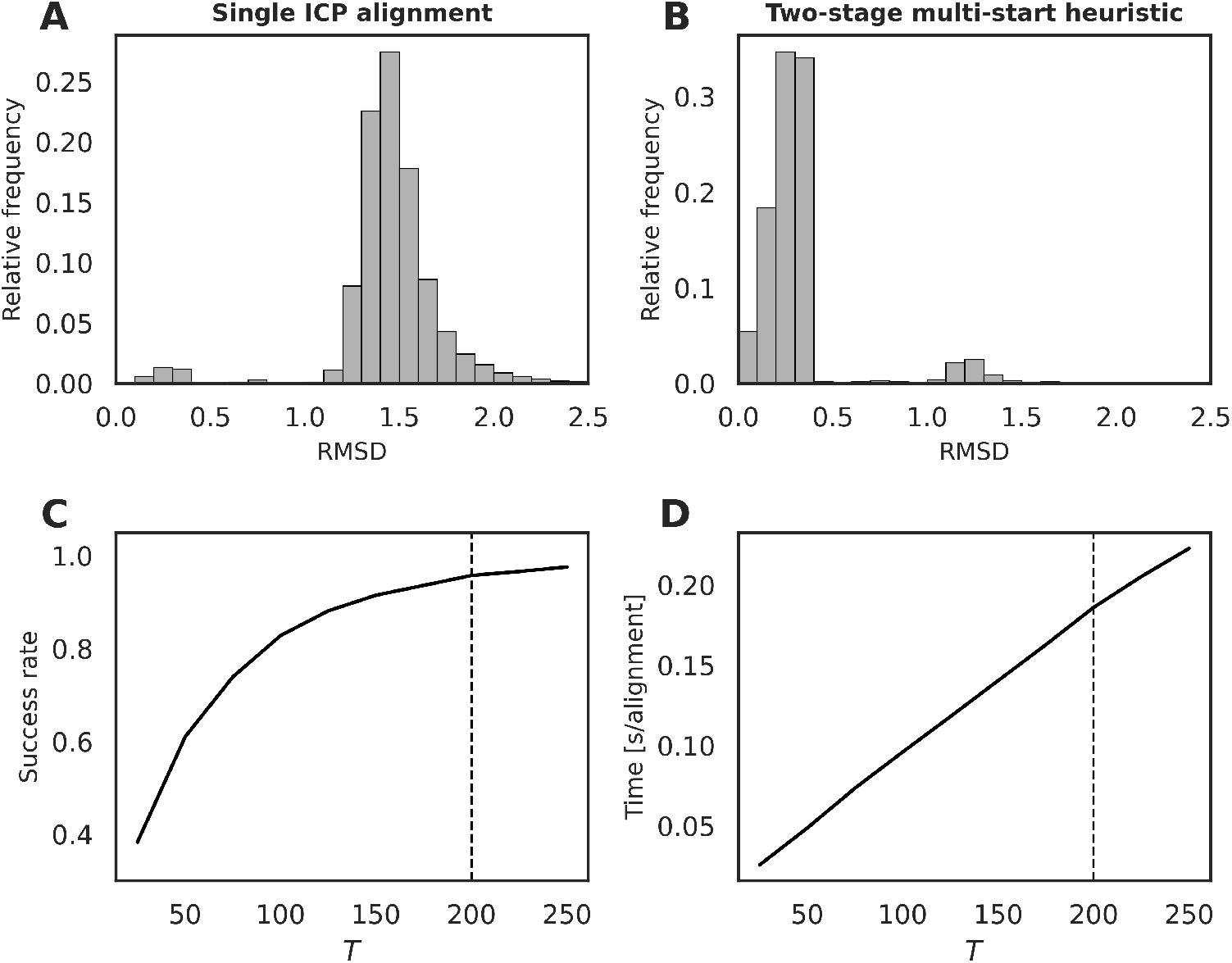
Performance of single-start and multi-start coarse-to-fine ICP alignment. Distributions of final RMSD values for 10^4^ geometrically similar MBS point-cloud pairs (reference minimum RMSD *<* 0.4 Å), aligned using either **A**. a single ICP run from one initialization or **B**. the two-stage, multi-start heuristic that screens *T* randomized coarse initializations and refines the top-scoring candidates. In this benchmark, a pair-specific reference minimum RMSD is defined as the lowest RMSD observed under an extensive multi-start protocol (*T*_*ref*_ = 500). **C**. Success rate, defined as the fraction of pairs whose heuristic result falls within a tolerance of the reference minimum, and **D**. wall-clock runtime per pair, shown as functions of the number of coarse initializations *T*. The dashed line indicates the selected operating point *T* = 200.

Although global point registration methods exist to generate ICP initializations (32), we found their performance to be inconsistent on these sparse, unordered point clouds, with a substantial fraction of alignments failing to reach the reference minimum achieved by repeated ICP (S2 Fig in S1 File). We therefore adopt a two-stage, multi-start coarse-to-fine ICP heuristic (Fig 1B) that screens a set of randomized initializations using a short ICP budget and then refines only the most promising candidates.

Before any restart, the point clouds are translated to a common origin by subtracting the centroid. All ICP runs in both stages use the same correspondence definition, correspondence-distance threshold, and robust loss settings as in the preceding subsection; only the iteration budget differs between stages. For each restart, the target point cloud is initialized by applying a random 3D rotation and then applying the fixed centering translation so that the initial translation is zero.

#### Two-stage heuristic

1. Course screening
  - Generate *T* independent randomized initializations as described above.
  - For each initialization, run ICP for *k* ≤ 5 iterations and compute an alignment score using the reported RMSD.
  - Rank-order the candidates by this coarse-stage RMSD score.
2. Refinement
  - Select the 5% highest-ranking candidates from the coarse screening.
  - Starting from each selected transform, run ICP with a larger number of iterations (*k* ≤ 30).
  - The final RMSD score is selected as the smallest of the resulting similarity metric results.

To quantify reliability, we benchmark the heuristic on 10^4^ MBS point-cloud pairs selected to have a low reference dissimilarity under repeated ICP alignment, thereby focusing on pairs for which a good alignment is attainable. For each pair, we define a reference minimum RMSD_*ref*_ as the lowest RMSD obtained across (*T*_*ref*_ = 500) randomized initializations using the same ICP objective and correspondence settings.

We count a heuristic run as successful if its final RMSD satisfies max(*ε* ∗ RMSD_*ref*_, 0.3) Å with *ε* = 1.2. We include this absolute floor 0.3 Å to avoid overly strict thresholds for pairs with very small RMSD_*ref*_, where minor numerical differences can dominate relative error. Using this definition, we report the success rate as the fraction of pairs meeting this criterion (Fig 1C). Balancing computational cost and convergence reliability, we select *T* = 200, yielding a 96% success rate with an average runtime of 0.19 seconds per alignment (Fig 1D).

### Representative MBS selection via RMSD-threshold clustering

Multiple PDB entries often capture the same protein in closely related conformations, and individual entries may provide several biological assemblies intended to reflect the functional oligomeric state. As a result, a single underlying MBS can appear multiple times in the extracted point-cloud dataset, inflating redundancy and increasing the likelihood of near-duplicate links in downstream network construction. To mitigate this effect, we apply a within-group redundancy reduction step to select a set of representative MBS point clouds prior to network assembly.

We first partition MBS point clouds into groups defined by shared UniProt accession and metal type. Within each group, we compute all pairwise MBS alignments using the two-stage ICP heuristic and obtain an RMSD score for each pair. We then construct a within-group similarity network in which nodes correspond to MBS point clouds and an undirected link is drawn between two nodes if their pairwise RMSD is below a fixed threshold (RMSD*<* 0.5 Å). Representative point clouds are selected using a greedy network-pruning procedure applied independently to each connected component of the within-group similarity network. For a given connected component, we compute for each node the mean RMSD to its adjacent neighbors and select the node with the smallest mean neighbor RMSD as the component representative. We retain this representative and remove all of its immediate neighbors (and their incident links) from the component, thereby eliminating near-duplicate point clouds within the threshold neighborhood of the representative. We repeat this select-and-prune step until no links remain in the component. All remaining isolated nodes are retained as representatives. The resulting set of retained point clouds across all groups constitutes the representative, redundancy-reduced dataset used for subsequent network construction.

### Network assembly from RMSD similarity with post hoc realignment

Given the redundancy-reduced MBS point-cloud set, we perform all-to-all pairwise alignments of the entire MBS dataset and form an undirected network in which each node corresponds to one MBS point cloud. A link is added between nodes *i* and *j* if their alignment score (i.e., RMSD) is below a global similarity threshold *τ*. We select *τ* in a data-driven manner by estimating how well low-RMSD outcomes separate from high-RMSD (misaligned) outcomes in empirical RMSD distributions. We stratify MBS pairs into bins defined by RMSD_*ref*_, and perform 10^3^ point cloud alignments to produce bin-wise empirical RMSD distributions. For each bin, we fit a two-component Gaussian mixture model and interpret the lower- and higher-RMSD components as representing geometry-consistent and misaligned or random-like outcomes, respectively. The two Gaussian components are defined as normal probability density functions

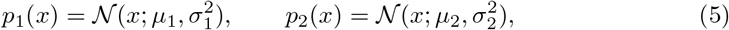

where 𝒩 (*x*; *µ, σ*^2^) denotes the Gaussian density with mean *µ* and variance *σ*^2^, and *µ*_1_, *σ*_1_ and *µ*_2_, *σ*_2_ are the means and standard deviations of the low- and high-RMSD components, respectively. To quantify the separability of the two components, we compute an overlap statistic *δ* defined as the area under the pointwise minimum of the two modes:

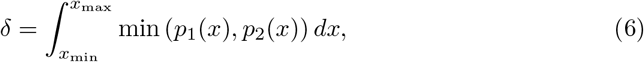

where the integration bounds are chosen to capture the effective support of both distributions:

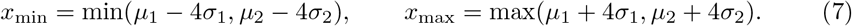

To define the global threshold *τ*, we track *δ* across bins where *δ* ≈ 0 indicates cleanly separated modes, whereas *δ >* 0 marks the onset of overlap, rendering the demarcation of well-aligned and misaligned pairs unreliable. We set *τ* conservatively to the upper end of the last bin preceding this overlap transition.

Because ICP outcomes remain initialization-sensitive even under the proposed multi-start heuristic, some true geometry-similar pairs may be missed by this primary thresholding step. To selectively revisit such candidate false negatives without re-aligning all non-links, we apply a topology-guided screening based on topological overlap (TO). We compute TO for node pairs that are not directly connected, defined as

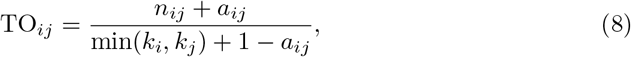

where *n*_*ij*_ is the number of common neighbors between node *i* and node *j, a*_*ij*_ denotes whether *i* and *j* are directly connected, and *k*_*i*_ and *k*_*j*_ are the degrees of nodes *i* and *j*, respectively. Pairs with TO ≥ 0.5 are re-aligned using the same ICP objective and correspondence settings but with an increased number of random initial transformations (*T* = 300) to further reduce the probability of convergence to a poor local minimum. If the resulting RMSD falls below the original threshold *τ*, we add a link between *i* and *j*.

### Metal ligand and functional co-occurrence

To quantify metal co-occurrence within the network, we test whether nodes annotated with metal *i* preferentially connect to nodes annotated with metal *j* relative to a permutation null. For each metal pair (*i, j*) among *M* metals, we compute the observed number of links *x*_*ij*_ whose incident nodes carry annotations *i* and *j*. We generate *R* = 10^4^ null networks by randomly permuting the node annotations while preserving the original network topology and degree sequence, and record the corresponding link counts 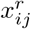. From these, we estimate *µ*_*ij*_ and *σ*_*ij*_ and compute the *Z*_*ij*_-score

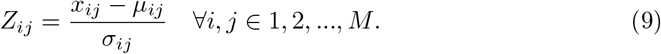

We also compute the two-sided empirical *p*-values

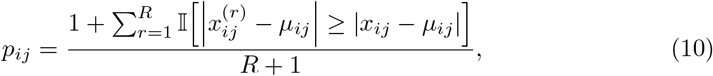

and adjust for multiple testing across all tested pairs (*i, j*) using the Benjamini–Hochberg procedure (33); pairs with FDR *<* 0.05 are considered significant.

We perform the same *Z*-score analysis for functional co-occurrence using the node’s EC numbers as categories, where the EC numbers are grouped at the third classification level.

### Computing network modules

To identify densely interconnected communities within components of the MBS similarity network, we perform community detection using the Leiden algorithm (34). Community partitions are computed on the weighted, undirected network, where link weights encode geometric similarity (i.e., RMSD scores) and thus emphasize stronger MBS–MBS similarity relationships. We used the *leidenalg* implementation (35) with resolution parameter *γ* = 1.

### Global structural alignment

We perform global structural alignments between protein structures using `TM-align` (36) via the RCSB PDB web interface (https://www.rcsb.org/alignment). Alignments are executed using default parameters, with individual PDB entries and chain identifiers specified as input. TM-scores are used as the primary metric for assessing global fold similarity, where values above 0.5 generally indicate similar global folds, and values below 0.3 are typically observed for unrelated structures (36).

### Joint analysis of local metal-binding site geometry and sequence context

We apply a two-stage sequence-similarity screening to protein-chains of connected MBS pairs in the network in order to characterize MBS similarity in the context of global and local sequence concordance. For each protein-chain pair, we compute an optimal global alignment using the Needleman-Wunsch algorithm (37) as implemented in Biopython (25) with the BLOSUM62 substitution matrix and affine gap penalties. Pairs with global identity *<* 25% are retained for local-homology screening.

For these pairs, we test for statistically significant local sequence similarity using Smith-Waterman local alignment computed with `ssearch36` (FASTA v36.1.1) (38). Alignments are run with composition-preserving shuffled-sequence controls enabled (-k 1000) and the BLOSUM62 matrix (`-s` BP62). Statistical significance is estimated from the shuffle-based null returned by `ssearch36`. To account for multiple testing and to express significance relative to the scale of our pairwise screen, we set the effective database size to the total number of local sequence alignments. MBS pairs with proteins exhibiting statistically significant local sequence similarity (E *<* 10^−3^) are excluded from further analysis.

To assess whether the geometry-conserved, sequence-divergent links identified by this screen are dispersed across the network or concentrated within locally dense regions, we compute the average clustering coefficient of the node-induced subgraph spanned by all MBS nodes incident to at least one retained candidate link. We evaluate this statistic against a permutation null by sampling, for *R* = 10^4^ replicates, link sets of identical size uniformly at random from the full MBS network, constructing the corresponding node-induced subgraph for each replicate, and recomputing its average clustering coefficient. A one-sided empirical p-value is then obtained as the fraction of null replicates with clustering coefficient greater than or equal to the observed value.

### Drug off-target inference from MBS-network enrichment and structural proximity evidence

This section describes the pipeline for identifying candidate drug off-target interactions supported by (i) enrichment of within-drug connectivity among known target MBS nodes in the MBS network and (ii) structural proximity evidence of drug binding near metal sites.

Drug-target interactions are obtained from DrugBank (v5.1.13). Targets are mapped to network nodes by UniProt accession, assigning each drug *d* the set of MBS nodes *V*_*d*_ corresponding to its annotated protein targets. Drugs associated with at least two MBS-containing target proteins are retained for analysis, designated as *interactor drugs*. If a target protein contributed multiple MBS nodes, all corresponding nodes are included in *V*_*d*_.

For each interactor drug, we quantify target interconnectivity as the number of links in the induced subnetwork over its target-node set, *x*_*d*_ = | *E*(*V*_*d*_) |. Statistical significance is assessed against a null distribution generated from *R* = 10^4^ degree-preserving randomizations of drug–MBS annotations. For each randomization *r*, we compute 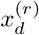, and derive a one-sided empirical *p*-value

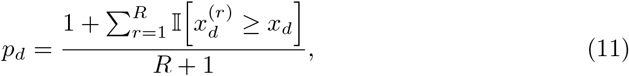

Empirical *p*-values are adjusted for multiple testing across all interactor drugs using the Benjamini–Hochberg procedure; drugs with FDR *<* 0.05 are considered connectivity-enriched.

For each connectivity-enriched interactor drug, we restrict the set of starting nodes to the subset of its target-associated MBS nodes that exhibit structural proximity evidence for that drug. Specifically, an MBS node is retained as a high-confidence starting point if the bound drug is observed within 5 Å of the node’s metal coordinate in the associated structure, or if the same proximity criterion is satisfied for any 1-hop neighbor of that node in the MBS network. Putative off-targets are then defined by expanding from these high-confidence starting nodes to their 1-hop neighbors that are not annotated as known targets of the drug in DrugBank.

### Software and data availability

All network construction and analyses are implemented in Python using NetworkX (39). Network visualizations are generated in Cytoscape (40) and accessed programmatically via its REST API using `py4cytoscape`. Source code for reproducing all analyses and figures is available at https://github.com/AlmaasLab/MBSNetwork.

## Results

### MBS dataset overview and impact of redundancy filtering

We assemble an initial metalloprotein dataset by parsing PDB entries containing any of the queried metals (Table 1) and extracting, for each metal ligand, an MBS point cloud from surrounding protein atoms (see Methods). Because closely related structures are frequently deposited multiple times (often with multiple biological assemblies), we apply a redundancy reduction procedure that clusters highly similar sites within protein- and metal-matched groups and retains representative point clouds. This filtering reduces the number of MBSs from 130,151 to 23,342, thus significantly decreasing structural redundancy, while retaining coverage of the same set of proteins (6,868). Key properties of both the retrieved and filtered datasets are summarized in Table 2.

**Table 2.** Summary of the properties of the retrieved and filtered MBS datasets.

|  | Retrieved dataset | Filtered Dataset |
| --- | --- | --- |
| MBSs | 130,151 | 23,342 |
| PDB entries | 25,846 | 12,142 |
| Proteins | 6,868 | 6,868 |
| MBSs with EC | 85,143 | 13,103 |

The filtered dataset largely preserves the relative metal and enzyme-class composition of the retrieved dataset, with a few notable exceptions (Fig 2). The most prominent change is a redistribution between Fe- and Zn-containing sites: Fe decreases from 39.8% of MBSs in the retrieved dataset to 26.0% after filtering, whereas Zn increases from 27.4% to 35.4% and becomes the most prevalent metal in the filtered dataset. The remaining metals show comparatively modest changes in relative abundance (e.g., Mn remains near 14%, and Cu remains near 8–9%). At the level of EC first-level classes, oxidoreductases (EC 1) remain the dominant class but decrease from 54.8% to 38.1% after filtering, accompanied by increases in transferases (EC 2; 9.8% to 16.0%) and hydrolases (EC 3; 24.4% to 33.8%).

**Fig 2.**
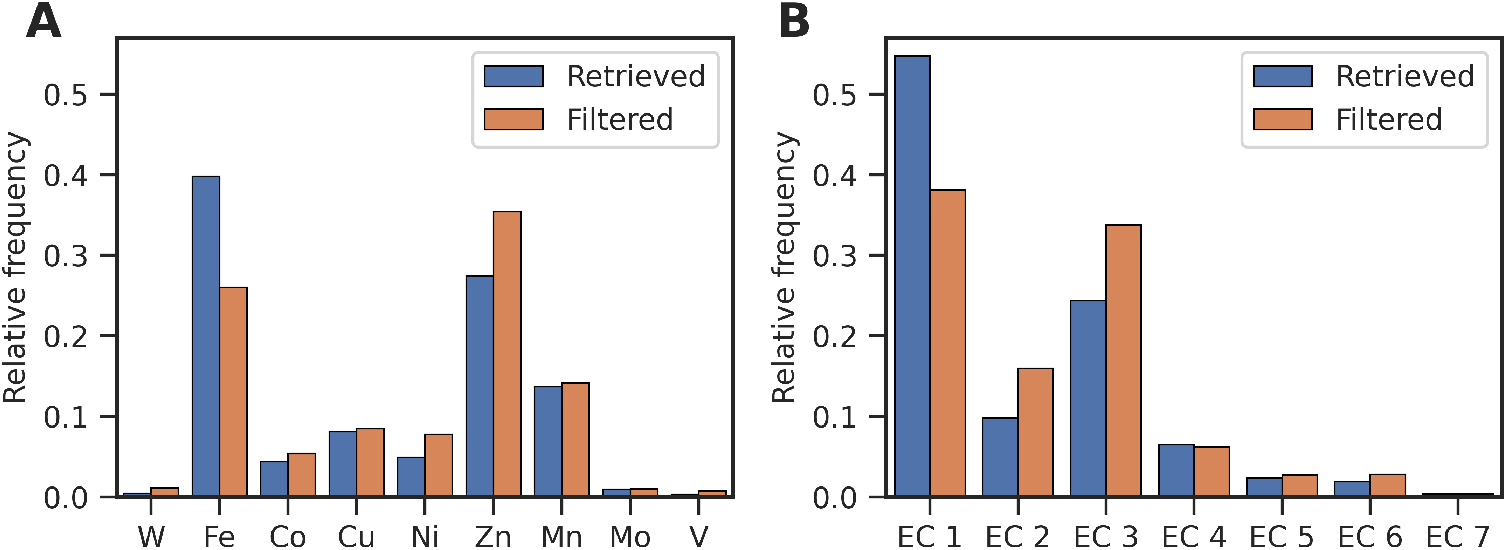
Distribution of metals and enzyme classes before and after redundancy filtering. Relative frequencies of **A**. metals and **B**. EC first-level classes (EC 1–6) in the retrieved dataset (all MBSs prior to redundancy filtering) and in the filtered dataset. EC-class frequencies are computed over MBSs with an assigned EC annotation.

### MBS network assembly with topology-guided link recovery

Using our two-stage point registration heuristic, we perform all-to-all pairwise alignment of the filtered MBS point cloud dataset (*N* = 23,342), corresponding to*N* (*N* − 1)/2 ≈ 2.7 × 10^8^ pairwise registrations and totaling approximately 14,250 CPU hours. We use the empirically observed bimodality of alignment RMSD values—reflecting separable regimes of good and poor alignment quality (Figs. 1A-B)—to guide selection of a conservative similarity threshold *τ* for network link formation.

To calibrate this threshold, we quantify the separation between the low- and high-RMSD modes across bins of increasing reference-minimum RMSD using the mode-overlap statistic *δ* (Eq. 6). As mode overlap increases, the distinction between the two regimes becomes less reliable. Based on the onset of appreciable overlap (Fig 3A), we select a global similarity cutoff of *τ* = 0.8 Å, corresponding to the upper end of the last bin preceding the overlap transition. The resulting global RMSD distribution from all pairwise alignments, with the selected cutoff indicated, is shown in Fig 3B. Applying *τ* yields a similarity network with 274,563 links (mean degree ≈ 23.5).

**Fig 3.**
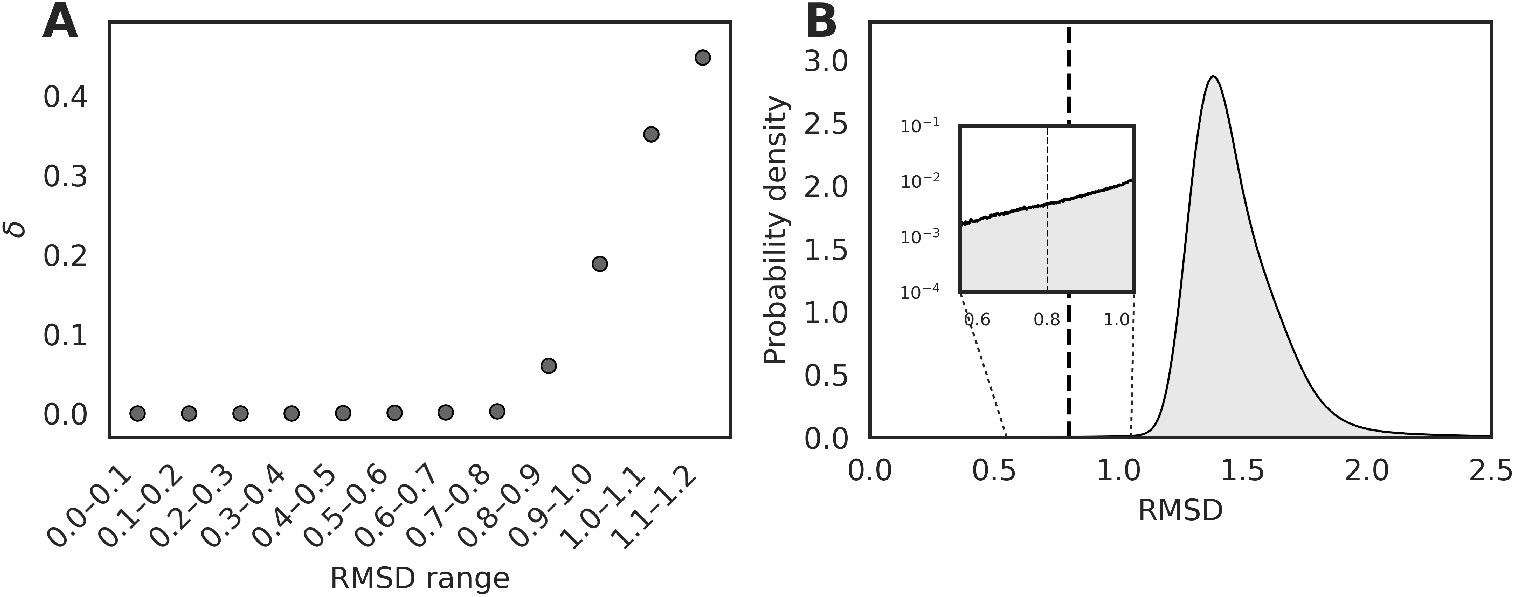
RMSD mode overlap and selection of the network similarity threshold. **A.** Mode-overlap metric *δ* estimated from two-component Gaussian mixture model fits to RMSD distributions computed from 10^3^ point cloud alignments per bin; bins are defined by increasing reference-minimum RMSD. **B**. Probability density of RMSD scores from all-to-all pairwise alignments of filtered MBS point clouds. The dashed line at *τ* = 0.8 Å marks the RMSD threshold used for network link creation.

As a targeted refinement to reduce potential false negatives due to the initialization sensitivity of ICP, we perform a post hoc realignment step on a restricted set of initially disconnected node pairs with TO ≥ 0.5 (Eq. 8). These candidates are re-aligned using an increased number of randomized initializations (*T* = 300). Pairs meeting the original cutoff after re-alignment are added as links, contributing an additional 37,510 links and increasing the network to 312,073 links (mean degree ≈ 26.7).

### Network connectivity reflects metal ligand type and functional co-occurrence

We next test whether connectivity in the MBS similarity network aligns with biologically interpretable node annotations, focusing on (i) metal identity and (ii) enzymatic function. If local geometric similarity is captured faithfully by the network links, we expect its topology to reflect both metal-specific coordination chemistry and shared functional properties of the associated proteins. Specifically, MBSs of the same metal should preferentially connect due to recurring coordination motifs, while MBSs from enzymes with related catalytic roles should exhibit enriched connectivity.

Visualizing the network at the selected similarity threshold (Fig 4) reveals a compartmentalized topology comprising 8,255 connected components. Despite this fragmentation, connectivity is locally dense (mean degree ⟨*k* ⟩ = 27; average clustering coefficient 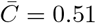), and overall modularity is high (*Q* = 0.92), consistent with MBS point clouds forming tightly connected neighborhoods separated by comparatively fewer between-neighborhood links. Inspection of the network further suggests that metal (Fig 4) and enzymatic function (S3 Fig in S1 File) composition appears structured across many components. This metal-based assortativity is particularly evident in the largest component *α* (Fig 4), in the fourth largest component, and across many of the intermediate-sized components, where links predominantly connect MBS nodes sharing the same metal ligand. In contrast, a few components, such as the second, third, and sixth largest component, display seemingly more heterogeneous compositions, with more frequent connections between MBSs of different metal types. Computing the relative distribution of metal-pair link composition across the entire network, we observe that for most metals the majority of links occur between nodes of the same metal (Fig 5A).

**Fig 4.**
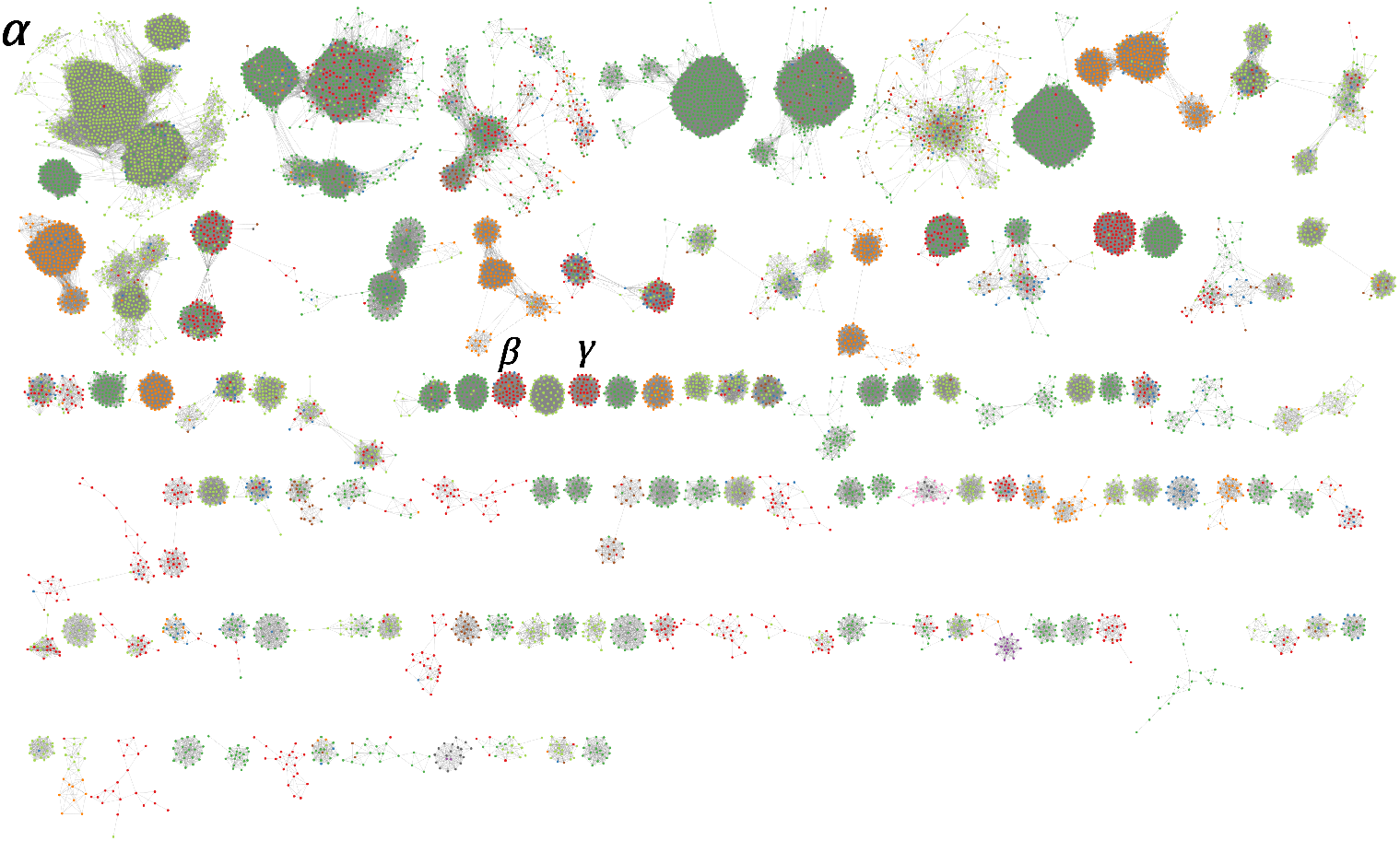
The MBS network colored by metal type. Connected components of size *N* ≥ 20 in the MBS network. Nodes are colored by the bound metal with the following coloring scheme: Co (blue), Cu (orange), Fe (dark green), Mn (red), Mo (purple), Ni (brown), V (pink), W (grey), and Zn (light green).

**Fig 5.**
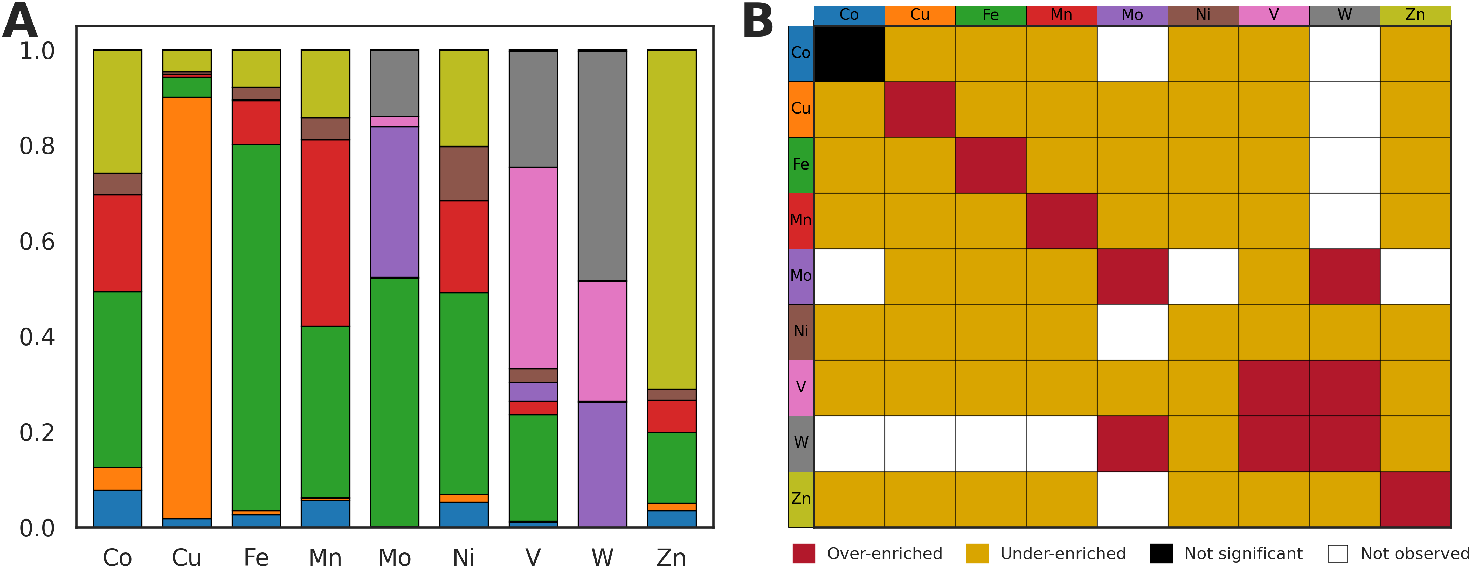
Metal-to-metal connectivity and enrichment. **A**. Observed relative distribution of metal-to-metal connectivity across network links. **B**. Heatmap of permutation-based *Z*-scores for metal-to-metal link counts, standardized against 10^4^ randomized label permutations. Significance is assessed using two-sided empirical *p*-values with Benjamini–Hochberg correction (FDR *<* 0.05). Significant enrichments are shown in color; non-significant cells are shown in black; metal pairs with no observed links are shown in white.

To rigorously quantify this pattern and account for differences in metal abundance and node degree that can inflate raw link fractions, we determine whether MBSs of the same metal are connected more frequently than expected by chance by performing a metal enrichment analysis. Specifically, we compute *Z*-scores for the number of links connecting nodes with metal *i* to those with metal *j*, using a null distribution generated from 10^4^ randomized networks. The resulting enrichment map (Fig 5B) is strongly diagonal, while most off-diagonal entries are significantly depleted relative to the null, indicating that network links largely connect MBSs of the same metal. In particular, Fe and Cu exhibit strong within-metal enrichment (Fe: *Z* = 120, *p* = 1.1 × 10^−4^; Cu *Z* = 134, *p* = 1.1 × 10^−4^), reflecting distinctive interaction profiles and suggesting limited structural compatibility with MBSs of other metal types. In contrast, between-metal enrichment is observed for W and V, and W and Mo. This is consistent with known cases where these metals can substitute in related enzyme families (41), but the network-based enrichment alone does not distinguish substitution from other sources of shared local geometry. Notably, Ni is the only metal to show a statistically significant within-metal depletion (*Z* = −3.3, *p* = 2.3 × 10^−3^), whereas Co displays no statistically significant enrichment (*Z* = 0.6, *p* = 5.4 × 10^−1^), hinting at broader geometric heterogeneity among their MBSs.

We next test whether network connectivity is similarly structured by enzymatic function. Using EC annotations grouped at the third level (a.b.c.*), we compute EC-pair link enrichment under the same topology-preserving label-permutation null used for metal enrichment and assess significance using two-sided empirical *p*-values with Benjamini–Hochberg correction (FDR *<* 0.05). The resulting *Z*-score heatmap (Fig 6) shows a prominent diagonal, indicating that MBS nodes assigned to the same EC subclass are connected more frequently than expected by chance. Compared to the metal-based analysis, the EC-based enrichment exhibits a larger number of significant off-diagonal associations, particularly among oxidoreductase (EC 1) subclasses. These cross-subclass associations suggest that some local metal-site geometries recur across distinct functional annotations, although attributing specific drivers (e.g., shared cofactor-binding motifs (42)) would require additional annotation of bound cofactors or domain context beyond the EC labels used here.

**Fig 6.**
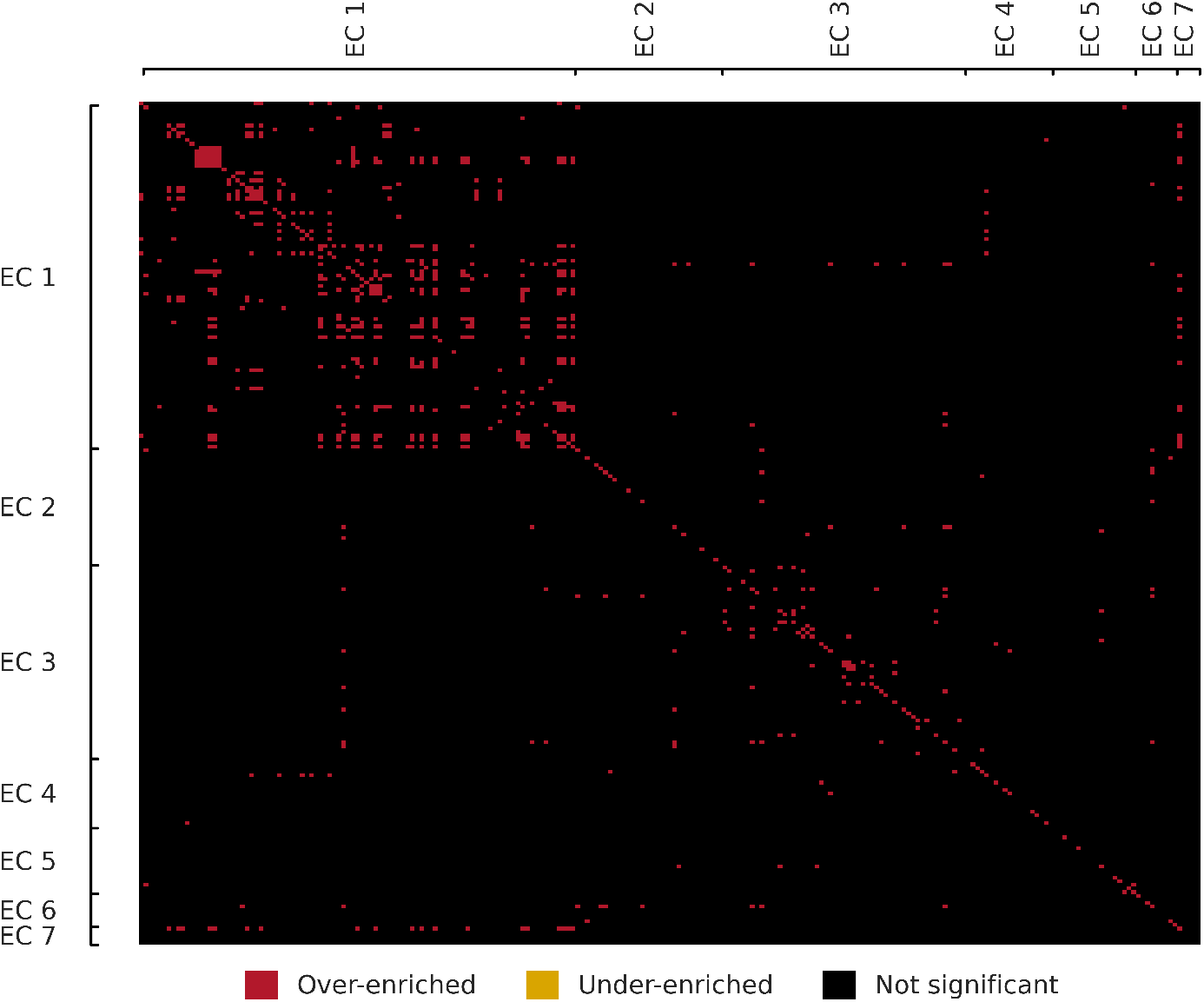
Enrichment of EC subclass co-occurrence among network links. Heatmap of permutation-based *Z*-scores for EC-pair link counts, with EC numbers grouped at the third level (a.b.c.*). Significance is assessed using two-sided empirical *p*-values followed by Benjamini–Hochberg correction (FDR *<* 0.05). Significant enrichments are shown in color; non-significant cells are shown in black.

### Recurrent motifs drive modularity of structural Zn-binding sites

The largest connected component of the MBS network (component *α* in Fig 4; *N* = 1,167 nodes) is enriched for Zn-binding sites. Many nodes in this component exhibit residue compositions characteristic of tetrahedral coordination environments dominated by cysteine and/or histidine, consistent with common structural Zn-binding architectures such as zinc-finger-like motifs (43; 44; 45).

To characterize internal structure within this component, we partition the component into modules using the Leiden community detection algorithm applied to the weighted network. We embed the component using a link-weighted spring-embedded layout, in which more similar MBS pairs exert stronger attractive forces (Fig 7), and summarize local residue environments by assigning each MBS a four-residue signature defined by the identities of the four residues closest to the metal coordinate (closest by minimum heavy-atom distance).

**Fig 7.**
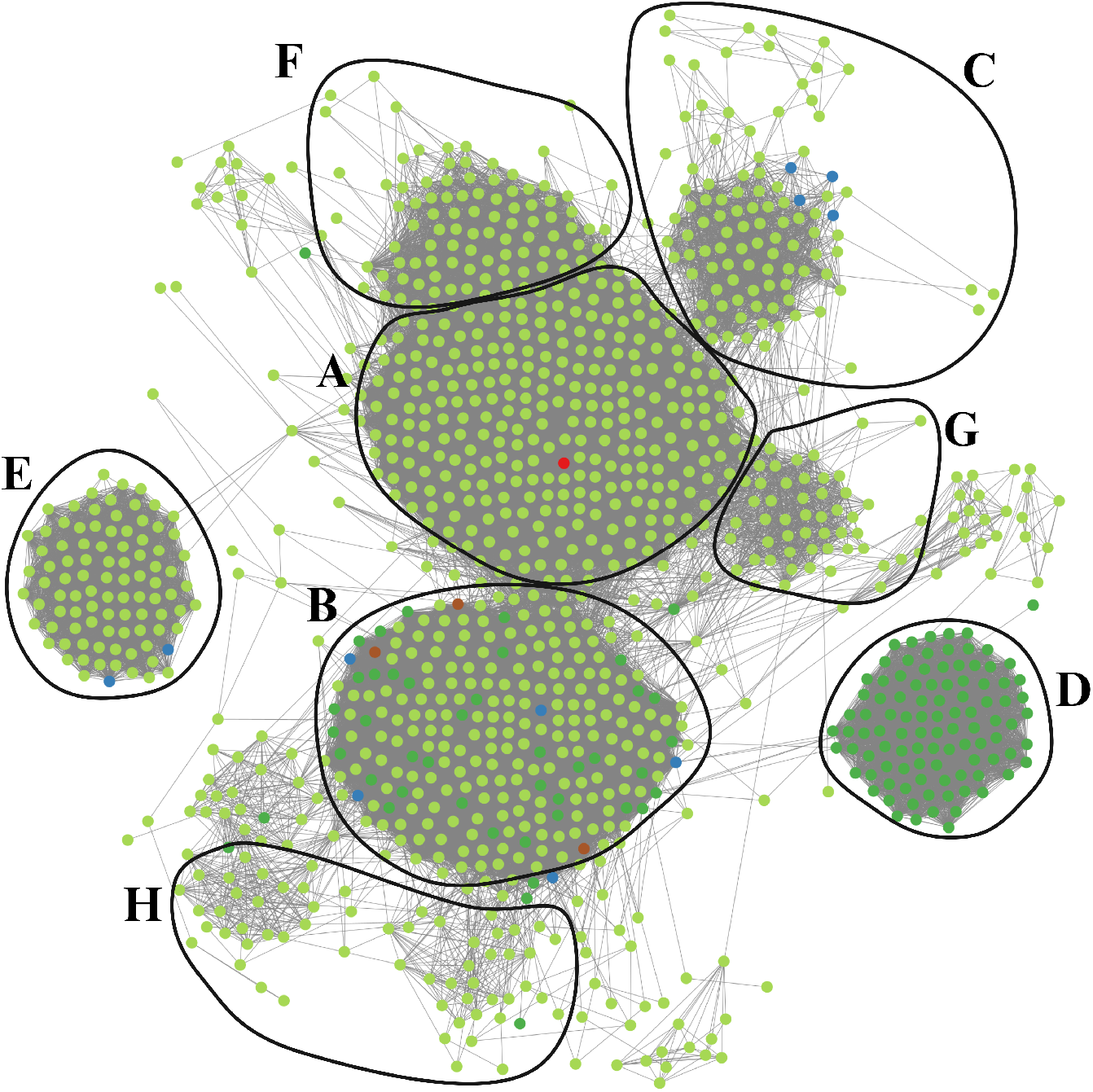
Modules of the largest connected component. Eight largest (*N* ≥ 50) Leiden modules within component *α* of the MBS network, visualized using a link-weighted spring-embedded layout (link weights proportional to geometric similarity). Nodes are colored by the bound metal ion: Co (blue), Fe (dark green), Mn (red), Ni (brown), and Zn (light green).

Focusing on the eight largest modules (A–H), each containing at least 50 nodes (Table 3), we find that four-residue signatures are strongly cysteine/histidine-enriched and align with canonical tetrahedral coordination patterns. Three modules (A, B, and E) are dominated by a Cys_4_ signature, with the dominant motif accounting for 74.6–91.4% of sites within each module, consistent with recurrent tetrahedral Cys-rich coordination environments. Three additional modules are enriched for mixed Cys/His coordination signatures (modules F, G, and H; Cys_3_His at 75.0%, 80.3%, and 52.5%, respectively), capturing common variants of Cys/His tetrahedral coordination. A further module (module C) is characterized by the canonical Cys_2_His_2_ motif, albeit with a lower dominant-motif frequency (56.0%), indicating greater within-module heterogeneity in residue composition despite tight geometric clustering. Metal identity is not perfectly aligned with module structure: module D is entirely Fe-associated (100%) yet exhibits a highly coherent Cys_2_His_2_ motif (91.8%), demonstrating that similar residue-level coordination architectures can place non-Zn sites within the same geometric neighborhoods as Zn-binding sites.

**Table 3.**
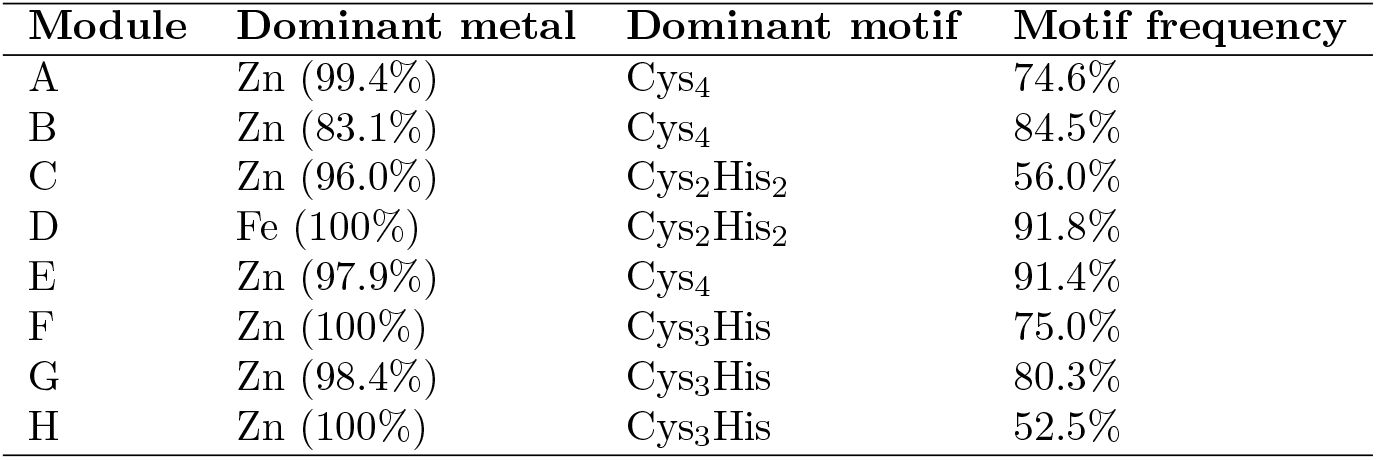
Coordination signatures of modules in the largest connected component. Each MBS is summarized by the identities of the four amino acid residues closest to the metal coordinate. Motifs are reported as unordered residue multisets (e.g., Cys_2_His_2_), with the dominant motif defined as the most frequent four-residue signature within the module.

These network components also include additional non-Zn sites embedded within otherwise Zn-enriched neighborhoods of the network. For example, module B contain several rubredoxin-like sites, in which an Fe ion is tetrahedrally coordinated by four cysteine residues (Cys_4_), a coordination geometry previously noted to resemble certain Zn-binding folds (46). Module D also contains sites consistent with Fe-S cluster coordination patterns that have previously been misassigned as Zn-binding in sequence-based analyses (47). More generally, the occurrence of non-Zn nodes within Zn-like modules supports the interpretation that recurrent coordination environments—captured here by local residue signatures and geometric similarity—can underlie module structure even when the annotated metal identity differs.

### Conserved metalloenzyme families form cohesive components in the MBS network

Given the enrichment of within-function connectivity observed at the EC-subclass level, we next ask whether this organization persists at the level of conserved metalloprotein families. Many metalloenzyme families retain highly similar local metal coordination environments across members, even when global sequence identity is low and overall fold context varies (48). If the MBS similarity network captures these conserved local architectures, then MBSs derived from a single family should preferentially connect, forming cohesive network structure.

As a case study, we examine the ureohydrolase domain superfamily, a well-characterized group of binuclear metalloenzymes with conserved active-site geometry and reaction chemistry (49). Members catalyze hydrolysis of non-peptide C–N bonds (EC 3.5.-.-) and employ a binuclear center in which two metals are coordinated by conserved aspartate and histidine residues within the active-site cleft of a three-layer *α*-*β*-*α* fold (50). We extract all ureohydrolase-associated MBS nodes in the network and examine their induced subnetworks, together with their immediate neighbors. Because each binuclear enzyme contributes two metal-centered MBS nodes (one per metal coordinate), we anticipate the selected ureohydrolase-associated MBS nodes to segregate into two components — one per metal-centered site in the binuclear catalytic center.

Consistent with these expectations, the ureohydrolase-associated MBS nodes partition into two distinct, highly connected network components (components *β* and *γ* in Fig 4) that correspond to the two metal-sites of the binuclear catalytic center (Fig 8). These components exhibit high internal connectivity and are functionally coherent, consisting primarily of MBSs (82 of 119) from arginase (EC 3.5.3.1) and agmatinase (EC 3.5.3.11), along with related ureohydrolase-domain enzymes such as formimidoylglutamase (EC 3.5.3.8) and N(*ω*)-hydroxy-L-arginine amidinohydrolase (EC 3.5.3.25). The components also display heterogeneity in metal identity, encompassing MBSs annotated with Mn, Ni, Cu, Co, and Zn within the same components, thereby providing another concrete example of between-metal connectivity in the network.

**Fig 8.**
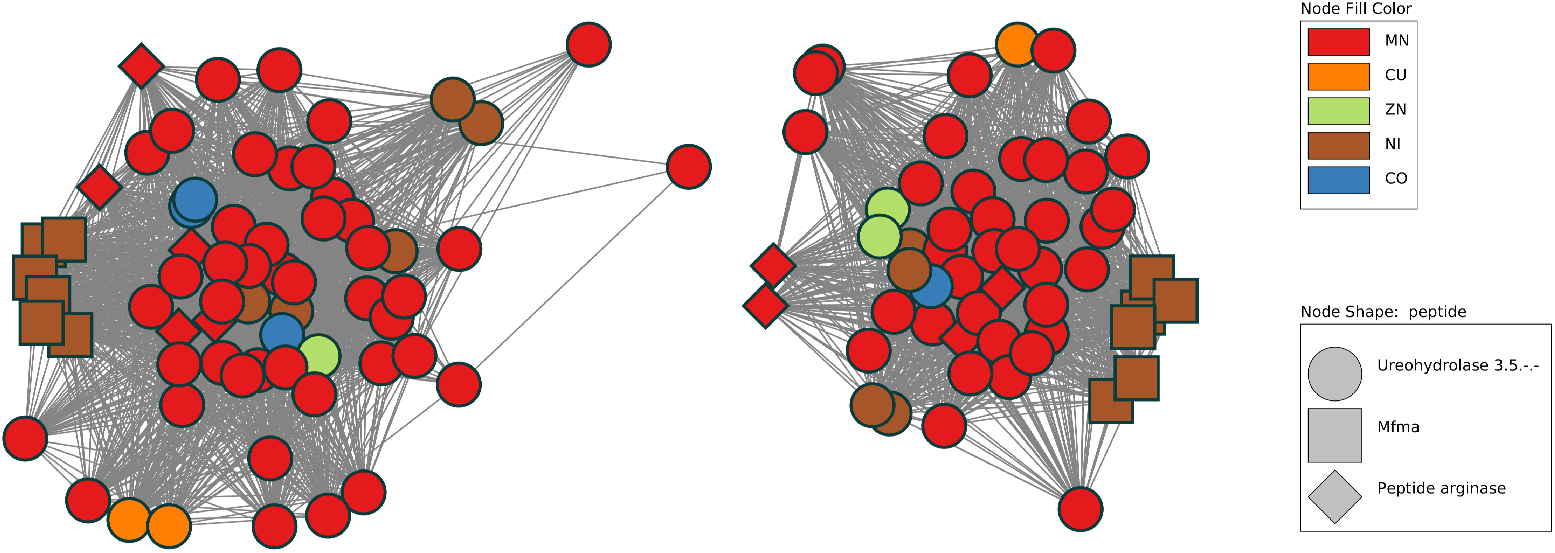
Ureohydrolase domain superfamily components in the MBS network. Shown are the two connected components formed by ureohydrolase-associated MBS nodes and immediate neighborhoods; each component corresponds to one metal-centered site of the binuclear catalytic center (i.e., one MBS node per coordinated metal ion). Node positions are computed using a force-directed layout with link weights derived from geometric similarity, so that more similar MBSs are placed closer together.

Beyond canonical ureohydrolases, these same components include atypical enzymes reported to preserve ureohydrolase-like binuclear site geometry. For example, MBSs from the recently characterized metformin hydrolase subunit A (*MfmA*) of *Ectopseudomonas mendocina* appear in both components, consistent with maintenance of a ureohydrolase-like local metal-site architecture despite only moderate sequence similarity to canonical ureohydrolases (51). A similar pattern is observed for the atypical peptide arginase from *Kamptonema* sp. implicated in post-translational arginine-to-ornithine modification: although this system differs in sequence context and quaternary organization (52), its MBS point clouds align closely with ureohydrolase-family MBSs and localize within the same two components. Together, these observations indicate that MBS-network connectivity can recover conserved local metal-site architectures within a metalloenzyme family, including relationships that are not readily implied by sequence- or fold-based similarity alone.

### Recurrent metal-binding site geometries across sequence-divergent proteins

As demonstrated, our point cloud alignment enable the identification of cases where highly similar MBS geometries occur in proteins with little sequence similarity. Another such example is provided in cluster 3 of component *α* (Fig 7). Within this cluster, an MBS of the phosphotyrosine-binding domain of the Hakai protein (CBLL1) contains an atypical Zn site in which a cysteine residue (Cys61) from the adjacent protomer completes the Cys_2_His_2_ coordination sphere (53). Aligning this site (PDB 3VK6) to a canonical Cys_2_His_2_ zinc finger (PDB 4M9E) yields a closely matching local geometry (RMSD 0.47 Å), despite low global sequence identity (20.1%) and poor global fold similarity (TM-score 0.22) (Fig 9). More generally, the occurrence of closely matching MBS geometries in proteins with little detectable sequence similarity is consistent with two non-exclusive scenarios: (i) divergent evolution from a remote common ancestor, in which the local MBS architecture is conserved while surrounding sequence and/or overall fold context diverges, or (ii) convergent evolution, in which similar coordination geometries and/or associated catalytic environments arise independently under shared physicochemical and functional constraints (54).

**Fig 9.**
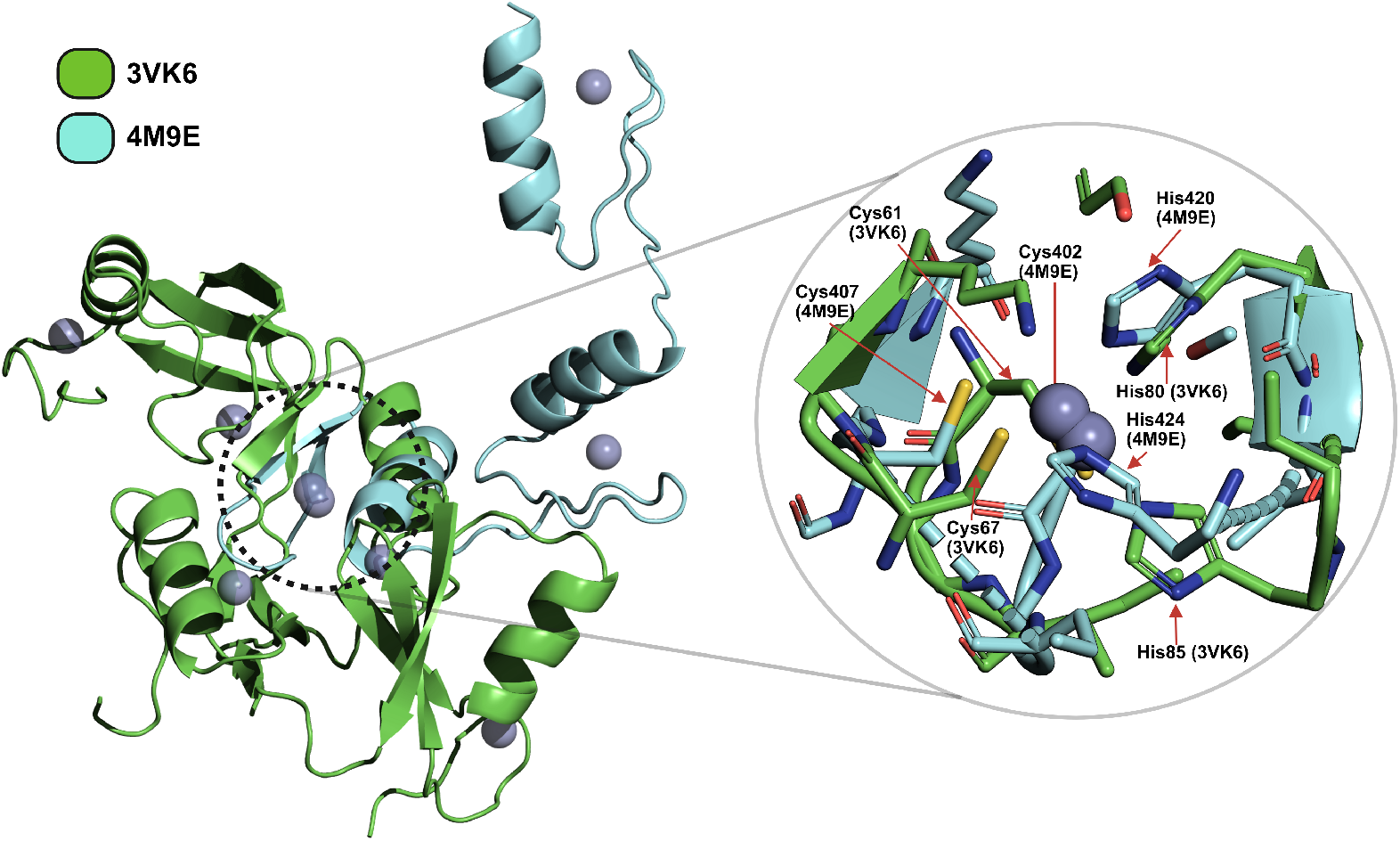
Structural alignment of Zn-binding sites from CBLL1 and a canonical Cys_2_His_2_ zinc finger. The Zn-binding site of the phosphotyrosine binding domain from E3 ubiquitin-protein ligase CBLL1 (PDB entry 3VK6, green) superimposed on a canonical Cys_2_His_2_ zinc finger domain from Krüppel-like factor 4 of *Mus musculus* (PDB entry 4M9E, cyan) using the optimal transformation obtained from the point cloud alignment (RMSD 0.47 Å). The inset show the protein structures that constitute the corresponding MBS point clouds and highlights the conserved tetrahedral coordination geometry of Zn (grey spheres).

To assess how frequently local geometry and sequence context decouple at network scale, we systematically compare protein sequence and MBS geometry across all network links. Specifically, we compute for each connected MBS pair in the network the global sequence identity of the corresponding proteins and compare these to the ICP-derived geometric similarity (max-normalized RMSD). Global sequence identity and local geometric similarity exhibit a moderate correlation (Spearman’s *ρ* = 0.37, *p* ≪ 10^−3^), indicating that higher sequence identity is generally associated with lower geometric divergence. However, inspection of the sequence–structure landscape reveals that this relationship is heterogeneous (Fig 10): the correlation is driven largely by MBS pairs with low sequence identity and only moderate geometric similarity, whereas strongly geometry-similar MBS pairs appear across a wide range of sequence identities. The latter includes regimes where sequence identity is minimal, motivating a targeted analysis of links where MBS geometry is conserved even at low sequence resemblance of the associated proteins.

**Fig 10.**
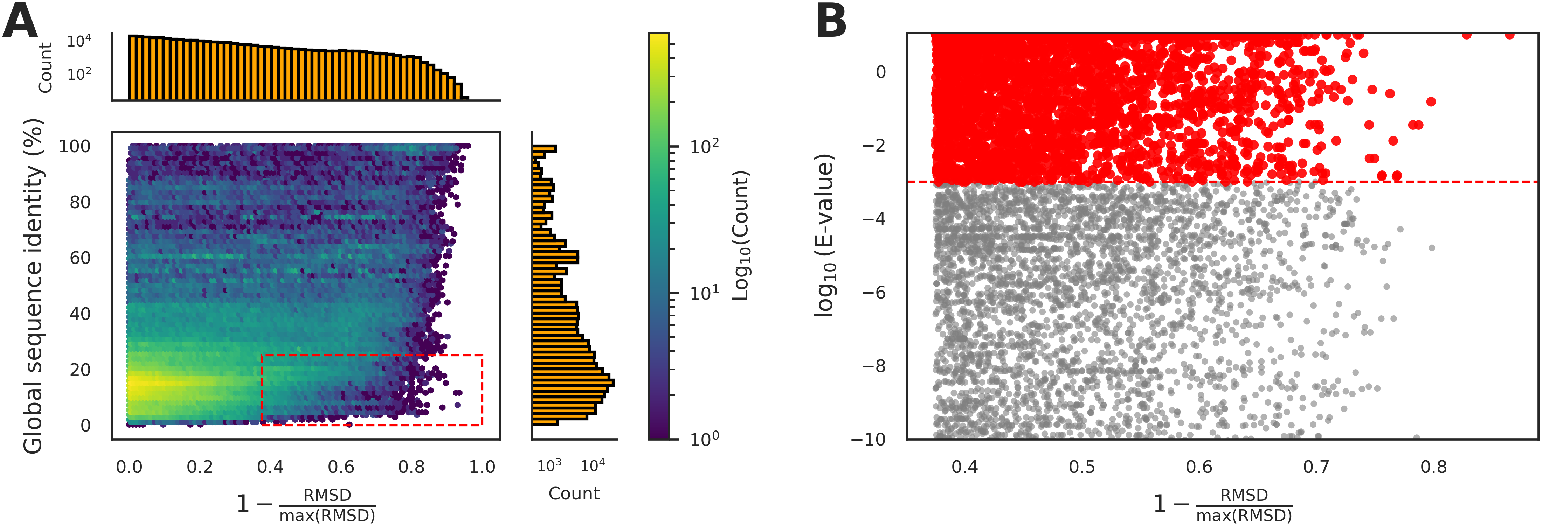
Sequence-geometry decoupling highlights recurrent metal-binding site geometries across sequence-divergent proteins. **(A)** Joint distribution of global sequence identity and normalized geometric similarity for all network-connected MBS pairs. The colorbar indicates the log-scaled density of MBS pairs in each bin. The red dashed box highlights MBS pairs combining low global sequence identity (*<* 25%) with high local geometric similarity (RMSD *<* 0.5 Å). **(B)** Local sequence similarity filtering for pairs in the red dashed box in (A). The log_10_(E-value) is obtained from shuffled-sequence Smith–Waterman alignments of the corresponding full-length proteins. The horizontal dashed line marks the threshold (*E <* 10^−3^) used to distinguish pairs exhibiting statistically significant local sequence similarity (grey points) from those without detectable local similarity (red points).

Accordingly, we compile MBS pairs that combine high geometric similarity with pronounced sequence divergence. We focus on network-connected MBS pairs (19,293) with low global sequence identity (*<* 25%) and high geometric similarity (RMSD *<* 0.5 Å) (red dashed box in Fig 10A). To enrich for cases without detectable local sequence homology, we further filter these pairs using pairwise local sequence alignment of the associated protein sequences; pairs with statistically significant local similarity at an expectation value threshold of *E* = 10^−3^ are excluded (Fig 10B). In total, this yield 7,237 MBS pairs spanning 1,159 unique proteins that exhibit high local geometric similarity with low global sequence identity and no detectable local sequence similarity under the chosen significance threshold. The full list of pairs, including their associated proteins, metals, and geometric and sequence similarity scores, is provided in S2 File.

Across the 7,237 candidate links, metal identity matches for most pairs (5,327; 74%), while 1,910 pairs (26%) connect sites annotated with different metals. The overall metal composition of the candidate set resembles that of the full dataset (S4 Fig in S1 File), indicating that geometry-conserved links across sequence-divergent proteins are not confined to a single metal chemistry. Functionally, 1,190 pairs share at least one EC annotation at the third classification level, thus constituting interesting cases of related catalytic activity despite lacking any appreciable sequence similarity under our filters. We further test whether the candidate links concentrate in locally cohesive regions of the MBS network by computing the average clustering coefficient of the node-induced subgraph spanned by their incident MBS nodes and comparing it to a permutation null generated by sampling link sets of identical size uniformly from the full network. The candidate-induced subnetwork shows a higher average clustering coefficient than expected under this null (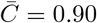, *p* = 10^−3^), indicating that these links are enriched within densely interconnected neighborhoods rather than being uniformly distributed across the network. Together, these candidate pairs delineate recurring metal-site architectures shared across sequence-unrelated proteins, providing a prioritized set for follow-up analyses with the aim of disentangling divergent evolution with local MBS conservation from structural and functional convergence.

### Network-driven drug off-target prediction

Misregulation of metalloproteins is a major contributor to human disease, and many drugs are designed specifically to target MBSs (55). Since drug–target interactions depend on structural compatibility, we posit that the topology of the MBS network can be leveraged to identify potential drug off-target interactions: If a drug is developed to bind to an MBS-containing protein, it may unintentionally bind to its target’s nearest MBS network neighbors owing to their similar MBS geometry. To investigate this possibility, we first map DrugBank drug-target interactions onto the MBS network, yielding 3,657 drugs with at least one MBS-containing target protein, collectively associated with 4,020 MBS nodes across the network. Among these, 800 drugs are linked to multiple MBS-encoding protein targets—hereafter referred to as *interactor drugs*—enabling a network-based assessment of whether a drug’s annotated targets occupy a coherent region of MBS space.

We next identify the set of MBS nodes targeted by the same interactor drug that are preferentially connected with each other in the network. Such topological enrichments indicate that multiple drug targets occupy closely related regions of MBS space, suggesting that the drug can engage a broader range of geometrically similar binding sites. When a drug’s MBS targets form these densely interconnected clusters, this pattern could also hint at reduced molecular specificity and an increased likelihood of cross-reactivity among structurally related metalloproteins. To delineate the subset of interactor drugs exhibiting significant target interconnectivity, we compute, for each drug, the number of network links among its target-associated MBS nodes and evaluate significance against a degree-preserving permutation null (see Methods). In total, we identify 326 interactor drugs with significantly enriched within-drug target connectivity after multiple-testing correction (FDR *<* 0.05; Fig 11), demonstrating that for a substantial subset of interactor drugs, the topology of the MBS network reflects known patterns of drug promiscuity.

**Fig 11.**
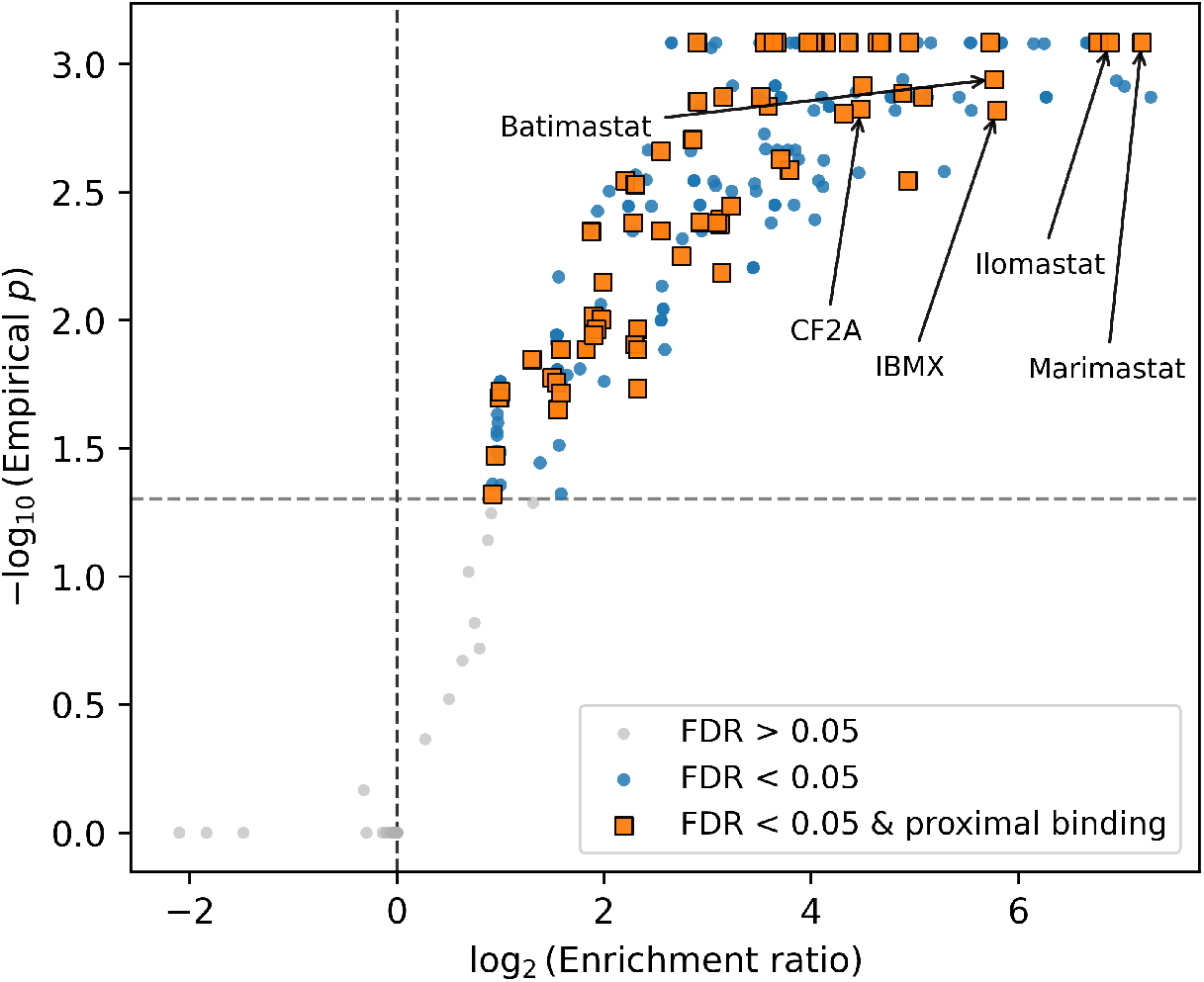
Network enrichment and structural proximity filters for drug off-target prediction. Volcano plot showing enrichment of interactor drugs, defined as compounds with ≥ 2 MBS–associated targets. The x-axis reports the enrichment ratio of within-drug target interconnectivity (number of network links among the drug’s target-associated MBS nodes) and the y-axis shows the Benjamini–Hochberg–adjusted empirical *p*-value obtained from the permutation test (*R* = 10^4^). Drugs passing FDR *<* 0.05 are considered connectivity-enriched. Square markers denote drugs with independent structural evidence of proximity, defined as at least one predicted binding pose located within 5 Å of a metal coordinate for at least one high-confidence starting node (or its 1-hop neighbor). IBMX, 3-isobutyl-1-methylxanthine; CF2A, 5-(2-chlorophenyl)-2-furoic acid.

To refine this signal toward higher-confidence off-target predictions, we next incorporate a complementary source of drug information. We define *proximal drugs* as those structurally observed to bind in the vicinity of an MBS, as curated from structural data (see Methods for details). This criterion confines enrichment to drug–MBS interactions that are consistent with metal-site proximity in available structures, rather than from coincidental binding to separate structural motifs that happen to co-occur with MBSs elsewhere in the protein. In total, we identify 1,233 proximal drugs associated with 6,363 MBS nodes in our dataset, and we focus on the subset of significantly enriched interactor drugs that also show proximal binding (135) near their associated MBSs. For these cases, geometric compatibility, as captured by the MBS network, is corroborated by independent experimental support of direct drug-to-MBS interaction (Fig 11). For each such drug, we infer putative off-targets by expanding from its high-confidence starting nodes to their 1-hop network neighbors that are not annotated as targets of that drug in DrugBank, restricting to human proteins to focus the analysis on clinically relevant off-target hypotheses.

Across the 88 connectivity-enriched drugs with structural proximity evidence, we recover 528 drug–off-target combinations spanning 151 unique human proteins (S3 File). For each predicted pair, S3 File reports the high-confidence starting node(s), the inferred off-target neighbor, and the intermediate network relationships used for inference. Several predictions correspond to drug–protein interactions previously reported in the literature but not annotated for the corresponding drug in DrugBank, providing case-level support that the MBS-network and proximity filters can recover independently observed off-target relationships. We therefore highlight a set of illustrative, high-ranking drugs for which multiple predicted off-targets have prior experimental support (Table 4).

**Table 4.** Selected network-prioritized candidate off-targets for connectivity-enriched drugs. Predicted human drug off-targets inferred from integration of MBS network topology and structural evidence of proximal drug binding. Predicted off-targets are categorized by level of supporting evidence: *Validated* — experimentally confirmed interactions reported in independent studies; *Inactive* — targets reported to exhibit no measurable activity against the drug; *Proposed* — targets hypothesized in prior literature based on indirect evidence (e.g., known target homologues); and *Predicted-only* — novel targets based on our network-driven approach.

| DrugBank | Drug | Predicted off-targets |
| --- | --- | --- |
| DB07954 | IBMX | MIOX, **PDE1B** (56), **PDE4C** (57), **PDE10A** (58), SAMHD1 |
| DB00786 | Marimastat | ADAM4, **ADAM8** (59), **ADAM17** (60), **ADAM33** (61), **ADAMTS1** (62), **ADAMTS4** (63), **ADAMTS5** (63), **BMP1** (64), ECE1, **TLL1** (65) |
| DB02255 | Ilomastat | ACE2, **ADAM8** (59), **ADAM17** (66), **ADAM33** (67), **ADAMTS1** (68), **ADAMTS4** (69), **ADAMTS5** (69), ANPEP, **BMP1** (70), **DPP3** (71), ECE1, ENPEP, **MME** (71), **MMP7** (72), **MMP10** (73), **MMP16** (74), **MMP20** (75), THOP1, TLL1 |
| DB03880 | Batimastat | **ADAM8** (59), **ADAM17** (76), **ADAM33** (77), **ADAMTS1** (78), **ADAMTS4** (79), BMP1, ECE1, **MMP1** (80), **MMP2** (80), **MMP7** (80), **MMP9** (80), MMP10, **MMP13** (81), **MMP14** (82), MMP20, TLL1 |
| DB02909 | CF2A | METAP1, METAP1D, METAP2, PEPD, XPNPEP1, XPNPEP3 |
|  |  | Validated<br>Inactive<br>Proposed<br>Predicted-only |

The prediction of 3-isobutyl-1-methylxanthine (IBMX) as an inhibitor of phosphodiesterase (e.g., PDE4C and PDE10A) aligns with experimental evidence of inhibitory activity (83; 57; 84). Likewise, among the predicted off-targets of the matrix metalloproteinase (MMP) inhibitors Marimastat, Ilomastat, and Batimastat, we recover multiple members of the ADAM and ADAMTS protein families, many of which have been proposed or experimentally validated as alternative targets (85; 61). Notably, the predicted off-targets for Ilomastat also include dipeptidyl peptidase III (DPP3) and neprilysin (MME), both having been identified as Ilomastat-sensitive metalloproteases by activity-based proteomic profiling (71). Known for their broad selectivity, these synthetic hydroxamate-type inhibitors have produced adverse effects during clinical trials, most notably musculoskeletal syndrome (MSS) (86). Although the mechanisms underlying MSS remain unresolved (87), the inhibition of non-MMP-members such as ADAM, ADAMTS, and related metalloproteases, including DPP3 and MME, represents a plausible contributing factor, consistent with their roles in extracellular matrix remodeling and inflammatory signaling (88).

Beyond cross-reactivity among human drug targets, our analysis also reveals potential off-target interactions for compounds originally developed against microbial protein targets. Notably, the antimicrobial agent 5-(2-chlorophenyl)-2-furoic acid (CF2A) is predicted to interact with several human metalloproteins, including type I methionine aminopeptidases METAP1 and METAP1D (Table 4). Given that CF2A was designed specifically to inhibit bacterial MetAP homologs, these findings suggest that its binding determinants are sufficiently conserved to permit interaction with the human isoenzymes. Such cross-species similarity highlights the importance of metal-site-aware similarity screening of metalloprotein-targeting antimicrobials, as inadvertent engagement of human metalloproteins could impact selectivity and safety profiles.

## Discussion

In this work, we extend and formalize the ICP-based framework of Raymond et al. (22) for aligning point cloud representations of MBSs. Whereas their study focused on a small set of dimanganese proteins and defined sites as all atoms within a fixed-radius sphere of the metal ion, we curate a comprehensive collection of all MBSs in the PDB and adopt a fixed-cardinality representation. Concretely, we define MBSs as a the *N* nearest protein atoms to the metal ligand based on an empirically calibrated estimate of site size inferred from local spatial neighborhoods around metal ligands of metalloprotein structures. This fixed-cardinality definition obviates the need for statistical techniques to interpret optimal alignments of differently-sized point clouds — which was deemed necessary due to the dataset size and *O*(*n*^2^) time complexity of all-to-all alignments. Despite its parsimonious nature, this representation proves remarkably powerful, yielding biologically meaningful geometric comparisons at scale.

A key challenge in adapting ICP for aligning MBS point clouds lies in its non-convexity, which makes the outcome strongly dependent on initialization and admits frequent convergence to suboptimal basins (32). In our setting, we observed alignment quality to be highly sensitive to the initial transformation. Although several methods have been proposed to improve initialization strategies (32), their poor convergence behavior (S2 Fig in S1 File), as well as computational cost, proved prohibitive for our approach. Our two-stage, multi-start coarse-to-fine heuristic addresses this by decreasing the likelihood of ending up with poor local minima while keeping the runtime manageable, and thus represents a pragmatic compromise between robustness and scalability. Importantly, while the heuristic reduces the incidence of false negatives — cases where structurally similar MBSs fail to be connected in the network — it cannot eliminate them entirely. To mitigate this, we implement a post hoc recovery step based on topological overlap to further recover structurally similar sites that have been missed due to unfavorable initializations. In contrast, the risk of false positives — spurious connections between structurally dissimilar MBSs — is controlled by the similarity threshold used to define network link formation. Since ICP registration lack a straightforward statistical interpretation (22), we derive this cutoff empirically by quantifying the separation between well-aligned and poorly aligned regimes in the bimodal RMSD distribution. This overlap analysis provides a principled rationale for our threshold selection, ensuring that links in the resulting MBS network reflect meaningful structural similarity rather than alignment artifacts.

The tendency of same-metal MBSs to connect preferentially in the network is consistent with both metalloprotein chemistry and the feature space induced by our point-cloud encoding. Many metals favor a limited repertoire of coordination numbers and geometries, and the first-shell arrangement of donor atoms and nearby backbone constraints can be highly conserved within common coordination motifs (6). Accordingly, metal identity explains a substantial fraction of network assortativity, as expected for an RMSD-based similarity network over local atomic neighborhoods. Importantly, however, the enrichment of connectivity within enzyme subclasses highlights that this signal extends beyond metal type alone. The local geometry of metalloenzyme active sites — defined by the precise configuration of coordinating residues and their immediate structural context — emerges as a key determinant of functional similarity, demonstrating how the very localized and exclusively geometric features of MBSs, as captured by our point cloud definition, are key determinants of both metal binding and biological function.

Consistent with this interpretation, we observe significant enrichment of connections within EC subclasses (Fig 6), supporting the view that conserved local geometries recur across functionally related metalloenzymes and that MBS alignment can be leveraged for functional inference. By embedding uncharacterized MBSs from newly determined structures into the network, their emergent neighborhoods may offer clues to functional roles. Such an approach could serve as a useful complement to sequence- and whole-structure annotation pipelines, particularly in the context of large-scale predictions from efforts such as AlphaFold (89) and AlphaFill (90), which are rapidly expanding the catalog of metalloprotein structures with unresolved functions. Because the pipeline operates on local MBS geometry alone, point clouds extracted from predicted structures can be embedded directly within the network to rapidly generate hypotheses about function and potential drug cross-reactivity for otherwise unannotated metalloproteins.

Nodes in our network represent individual metal-binding sites, whereas functional metadata (e.g., EC numbers) are assigned at the protein-chain level. Consequently, an MBS can inherit the catalytic annotation of its host enzyme even when the specific metal site is auxiliary—structural, regulatory, or distal from the catalytic center. This label mismatch is exemplified by module 5 in component *α* (Fig 7), which includes numerous MBSs from Zn-containing alcohol dehydrogenases (ADHs, EC 1.1.1.1): the assigned EC number reflects the enzyme’s catalytic function, yet the MBSs correspond to structural Zn-binding sites that are distinct from the catalytic Zn site (91). A similar ambiguity arises in 3-hydroxyanthranilate 3,4-dioxygenase (EC 1.13.11.6), where annotated metal sites have been proposed as structural rather than catalytic (92). These examples emphasize an important caveat: While the MBS network reliably captures local geometric similarity, the functional interpretation of individual sites requires care, as not all annotated MBSs correspond to sites of catalytic activity. Nevertheless, the inclusion of auxiliary MBSs does not undermine the validity of functional inference. These sites often contribute to protein stability, cofactor positioning, or assembly interfaces, roles that can be highly conserved across functionally related enzymes. Thus, their recurrence within the network still encodes biologically meaningful information, broadening the scope of functional inference beyond enzyme catalysis alone.

Using network connectivity to integrate local geometric similarity of MBSs with sequence similarity of the encoding proteins, we systematically examine how conservation at the level of local geometry relates to evolutionary relatedness at the sequence level. This joint analysis indicates that, although global sequence identity and MBS geometry exhibit a modest correlation, this relationship is highly non-uniform across the sequence–structure landscape (Fig, 10). In particular, the correlation is largely driven by pairs with low sequence identity and moderate geometric similarity, whereas among the most geometrically similar MBSs the corresponding sequence identities vary widely and show no clear dependence on local structural similarity.

The occurrence of highly similar MBS geometries in proteins that are otherwise strongly divergent in sequence admits two evolutionary interpretations. In some cases, they may reflect very remote common ancestry, in which the metal-binding core is selectively retained while the remainder of the protein diverges (93). In this view, metal-binding motifs can act as deeply conserved functional elements that persist over long evolutionary timescales even as global sequence similarity is eroded (6). Alternatively, closely matching coordination environments may arise through the repeated selection of similar structural solutions under comparable physicochemical and functional constraints, whereby unrelated protein scaffolds independently adopt analogous MBSs to support similar biological functions (94). In this context, the MBS pairs identified here provide a systematic collection of examples that can be used to further delineate and help distinguish deep evolutionary MBS conservation in otherwise divergent protein sequence contexts or the convergent emergence of MBSs across unrelated protein lineages.

We combine MBS-network topology with experimental evidence of drug proximity to identify metalloproteins that are plausibly susceptible to shared drug interactions. By focusing on drugs whose protein targets’ MBSs exhibit significant network connectivity and confirmed proximal binding to the MBS, we delineate a high-confidence set of drug–off-target pairs spanning a diverse range of human metalloproteins (S3 File). By integrating experimental binding evidence with network topology and connectivity enrichment, our framework extends drug annotations beyond established targets, capturing experimentally verified interactions overlooked by existing resources and novel candidates. The framework’s recovery of experimentally validated but under-annotated off-targets demonstrates its capacity to detect biologically relevant drug–protein interactions. Notably, the accurate identification of DPP3 and MME as Ilomastat off-targets — interactions independently confirmed through activity-based proteomic profiling (71) — provides direct empirical validation of the predictive fidelity of our approach. Although MME shows no discernible sequence homology to MMPs (71), its recovery as an Ilomastat off-target underscores how shared MBS geometries can reveal functional relationships invisible to sequence-based analyses. These examples demonstrate that the MBS network captures the underlying physicochemical determinants of inhibitor binding, such as conserved metal coordination geometry and pocket topology, which may transcend sequence or fold similarity. The consistent agreement between our predictions and experimentally confirmed off-targets of hydroxamate-based metalloproteinase inhibitors further indicates that the network topology effectively encodes structural logic of drug promiscuity. Collectively, these findings substantiate the framework as a scalable and mechanistically grounded approach for off-target inference, offering a means to identify unrecognized cross-reactivities with direct translational relevance to drug selectivity optimization, adverse effect assessments, and the broader study of polypharmacology.

While our work here centers on MBSs, the broader idea — encoding localized protein microenvironments as point clouds and comparing them with rigid point cloud registration — could be extended to other functional regions, including non-metal enzyme active sites. Constructing similarity networks from such representations would naturally support motif retrieval and clustering, enabling the prioritization of candidate catalytic pockets in newly solved or predicted protein structures by highlighting recurring geometric patterns across enzymes with related chemistry. More generally, local site representations can serve as inputs to graph-based models for predicting enzymatic properties, including kinetic parameters such as *k*_cat_ and *K*_M_. Recent work has shown that graph neural networks operating on full protein structures can predict kinetic constants (95; 96); however, global representations risk diluting the functional signal with structural features irrelevant to catalysis. From a machine learning perspective, it represents a poor signal-to-noise trade-off: The model is tasked with parsing the entire fold architecture when the information most predictive of enzymatic function is concentrated in a small fraction of the structure. We therefore expect that models explicitly centered on active-site microenvironments will offer a more direct route to learning structure–kinetics relationships. Consistent with this view, geometric descriptors of active sites have been shown to carry predictive signal for enzymatic turnover rates (97), motivating future extensions of local-structure similarity networks beyond metalloproteins to broader questions of catalytic function and quantitative enzymology.

## Acknowledgments

The authors would like to express their gratitude to Emil Karlsen for his assistance with the setup and execution of the all-to-all pairwise point cloud alignment on our in-house compute cluster. We also thank Gaston Courtade for constructive and valuable feedback on the manuscript.

## Supporting information

**S1 File** Supporting figures.

**S2 File** MBS pairs with high geometric similarity coupled with low sequence similarity between their encoding proteins, together with associated protein-level annotations and metadata.

**S3 File** Predicted human drug off-targets identified by integrating MBS network connectivity enrichment with structural proximity evidence.

