## Supplementary material for "Metal binding site alignment enables network-driven discovery of recurrent geometries across sequence-divergent proteins and drug off-targets": S1 File

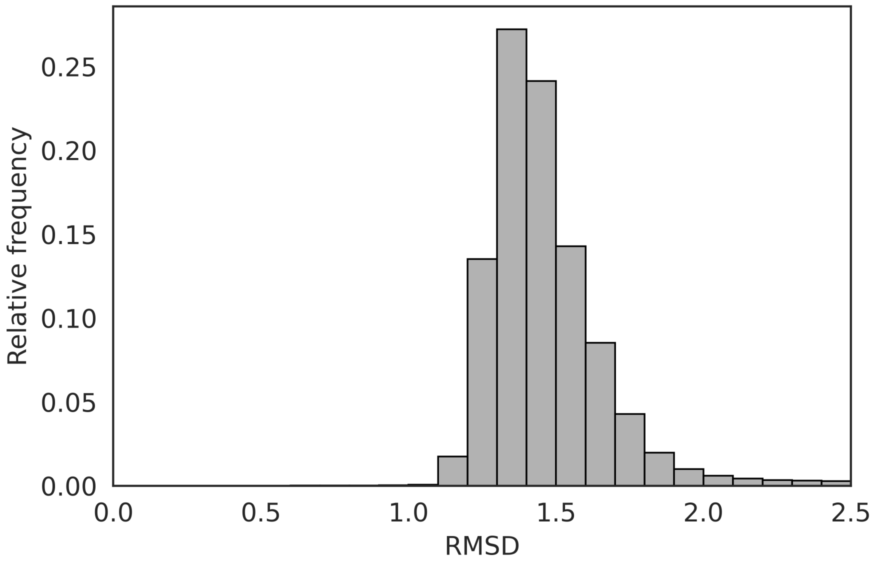


**Fig S1:** Distribution of RMSD values computed for $10^5$ randomly selected MBS point cloud pairs, providing a baseline which characterizes the expected RMSD distribution when aligning unrelated structures.

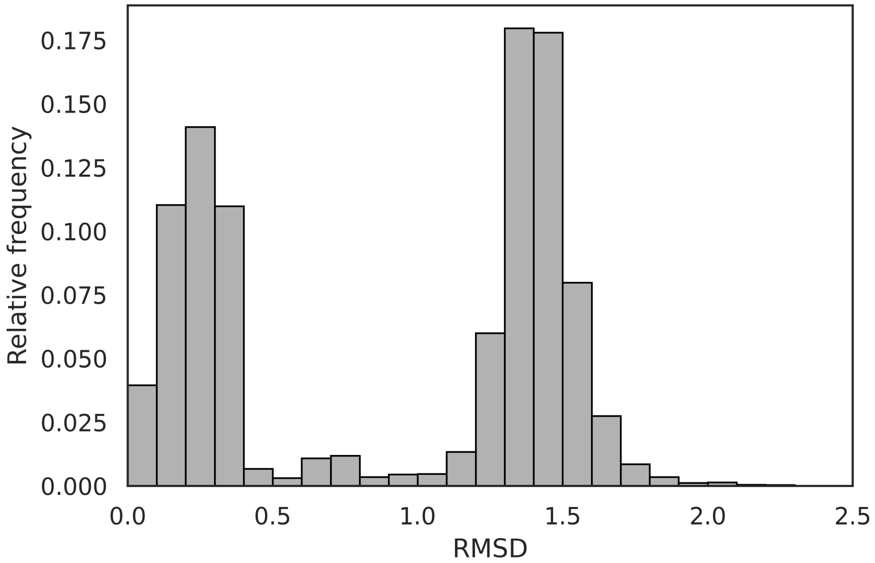


**Fig S2:** Distribution of RMSD values for ICP alignment of $10^4$ pairs of structurally similar MBS point clouds (global-optimum RMSD $< 0.4$ Å). Initializations were generated using Fast Global Registration, followed by fine-tuning using ICP.

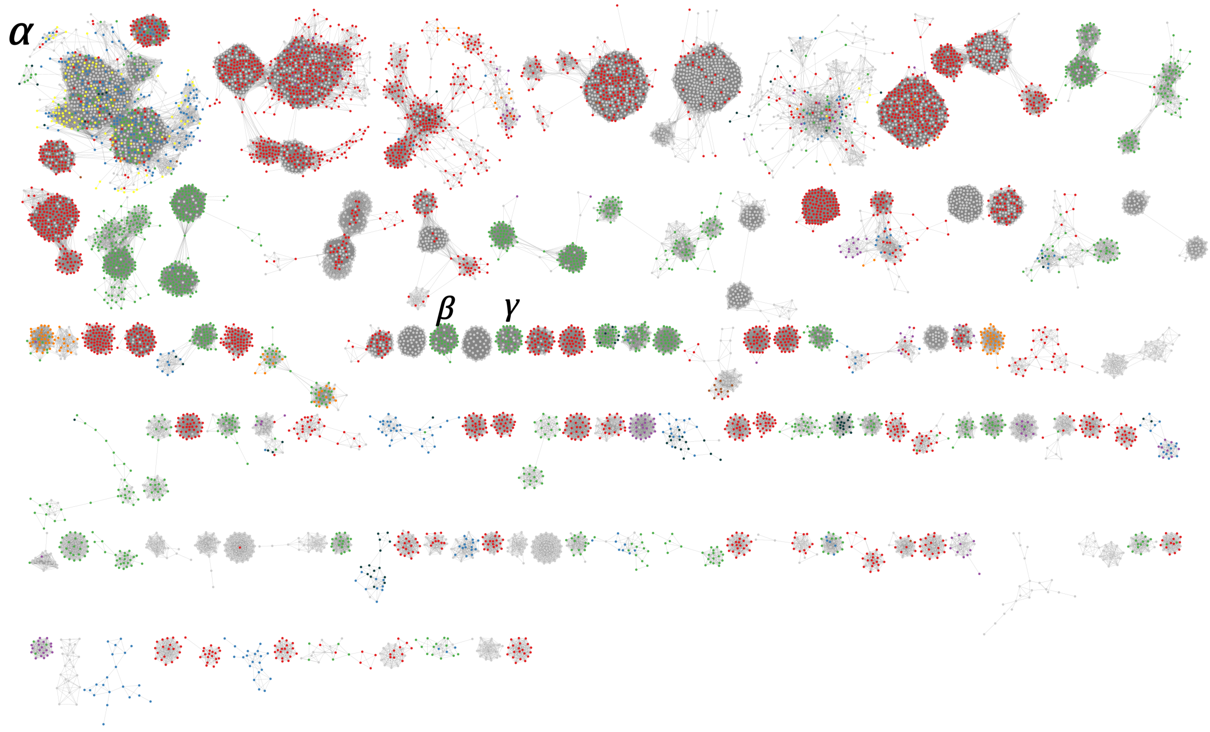


**Fig S3:** Connected components of size $N \geq 20$ of the MBS network. The nodes are colored according to the highest classification digit of their Enzyme Commission (EC) number. Oxidoreductases (EC 1, red), transferases (EC 2, blue), hydrolases (EC 3, green), lyases (EC 4, purple), isomerases (EC 5, orange), ligases (EC 6, yellow), translocases (EC 7, brown). Nodes with ambiguous or undefined EC assignment are colored black and grey, respectively.

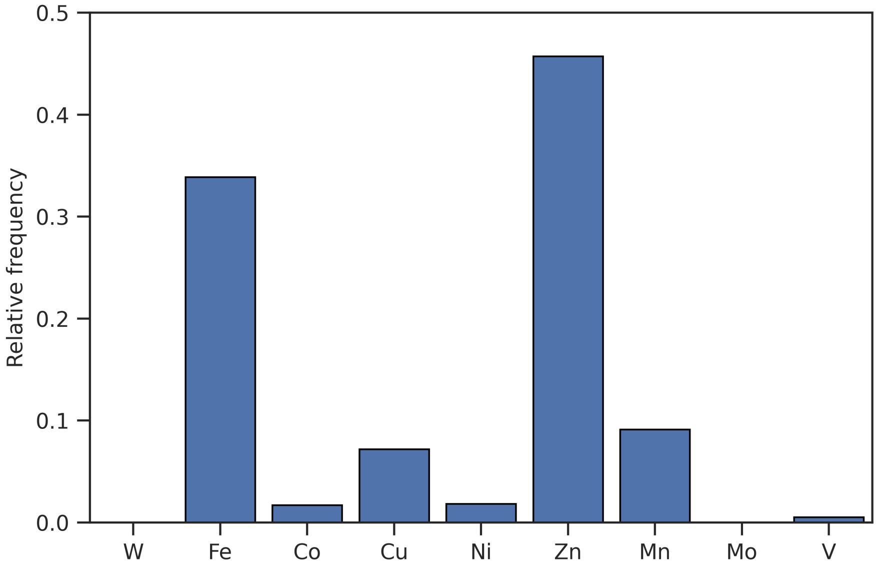


**Fig S4:** Relative distribution of metal ligands of the set of identified MBS pairs with highly similar point cloud geometries across sequence-divergent protein contexts.
